# REVOLUTA regulates cell fate and wall patterning in the fruit endocarp

**DOI:** 10.64898/2026.04.20.719616

**Authors:** Aurélia Emonet, Ilsa Spatz, Ciaran Kelly, Nikita Tikhomirov, Sonja Dunemann, Lisa Brombach, Michael Raissig, Markus Pauly, Angela Hay

## Abstract

Explosive seed dispersal distinguishes *Cardamine* species from Arabidopsis and depends on polarized secondary cell wall (SCW) deposition in fruit endocarp *b* (end*b*) cells. How this SCW pattern is specified and environmentally modulated remains unclear. The polyploid *Cardamine chenopodiifolia* produces explosive aerial fruit and non-explosive subterranean fruit, creating a tractable system to address this problem. We show light triggers underground fruit to explode by reprogramming end*b* SCW patterning from uniform to polar. We identify the HD-ZIPIII transcription factor REVOLUTA as a central regulator of end*b* cell fate, SCW formation, and organ polarity in Arabidopsis and *Cardamine hirsuta*. In *C. hirsuta*, duplicated *REVOLUTA* paralogs are required for end*b* SCW deposition, while other *HD-ZIPIII* genes contribute redundantly to cell fate and organ polarity. *REVOLUTA* over-expression converts polar end*b* SCWs to uniform, producing non-explosive fruit. Together, these findings reveal a tunable developmental module underlying evolutionary transitions between explosive and non-explosive seed dispersal strategies.

## Introduction

Understanding how complex traits develop and diversify is a central question in biology. Fruit diversity reflects the wide range of strategies plants use for seed dispersal^1^, a process with important ecological and evolutionary consequences^2,3^. A striking example of this diversity is the exploding seed pods of *Cardamine* (Brassicaceae)^4^, a trait that sets this genus apart from the closely related model plant Arabidopsis. The small weed *Cardamine hirsuta* achieves ballistic seed dispersal through ultrafast coiling of its fruit valves, launching seeds at speeds exceeding 10 m/s^5^. This complex biomechanical trait results from coordinated innovations operating across multiple spatial scales^5^.

At the organ scale, elastic energy stored in the fruit valves powers explosion. This energy results from tension generated by differential contraction of valve tissues^6^, and its rapid release is controlled at the cellular scale by the geometry of lignified secondary cell walls (SCW) in endocarp *b* (end*b*) cells located on the inner adaxial side of the fruit valve^5^. In non-explosive Arabidopsis fruit, end*b* SCWs are deposited uniformly^7^, whereas in explosive *C. hirsuta* fruit they adopt a distinctive polar pattern. Mathematical modeling demonstrated that this polar SCW pattern enables fruit valves to coil rapidly, analogous to a toy slap bracelet^5^. Genetic and phylogenetic analyses further established that polar end*b* SCW patterning is both required for and strictly associated with explosive seed dispersal in *Cardamine*^5^. Together, these findings identify polar SCW patterning in end*b* cells as a key innovation underpinning the evolution of explosive seed dispersal.

The genetic mechanisms that specify end*b* cell fate and SCW patterning, and how they evolved to generate diverse seed dispersal traits, remain largely unknown. In Arabidopsis fruit, SCW formation is controlled by the transcription factors NAC SECONDARY WALL THICKENING PROMOTING FACTOR 1 and 3 (NST1/3)^8^. NST1/3 do not specify end*b* cell identity, instead they activate a regulatory cascade that drives SCW synthesis in end*b* cells^8–10^. In *C. hirsuta*, end*b* SCW formation requires the assembly of CESA7-containing cellulose synthase complexes at the plasma membrane^11^, as well as three secreted laccases (LAC4, 11 and 17) that spatially pattern lignin deposition within the cell wall matrix^12^. Although these laccases are necessary for polar lignin deposition in *C. hirsuta* end*b* cells, they are not sufficient to impose polar patterning when transferred to Arabidopsis^12^. Thus, it remains unclear whether polar end*b* SCW patterning alone is sufficient to confer explosive seed dispersal to non-explosive fruit.

To explore this question from a new perspective, we developed experimental tools in the amphicarpic species *Cardamine chenopodiifolia*^13,14^. This species produces two morphologically distinct fruit types on the same plant—one above ground and the other below ground. Aerial fruits retain polar end*b* SCWs and explode like other *Cardamine* species. In contrast, subterranean fruits exhibit uniform SCWs and are non-explosive, resembling Arabidopsis^13^. This dual seed dispersal strategy constitutes a bet-hedging mechanism that can be advantageous in unpredictable environments^15^. Comparing transcriptomes of aerial and subterranean fruit valves revealed fruit type-specific expression of cell wall-related genes, but did not pinpoint regulators of end*b* cell fate or SCW patterning^13^.

The emergence of an evolutionary novelty such as amphicarpy depends on both the genetic and environmental context. In *C. chenopodiifolia*, divergent fruit forms are produced from a single genome but develop in sharply contrasting environments above and below ground. Studying this trait therefore provides a powerful opportunity to dissect how environmental cues interact with developmental programs to generate phenotypic diversity^16–18^. Consistent with this idea, fruit valve transcriptomes in *C. chenopodiifolia* exhibit strong environmental signatures^13^. Aerial fruit are characterized by up-regulation of photosynthetic and light-responsive genes, whereas subterranean fruit—deprived of light and exposed to the soil microbiome—show enrichment of defense response pathways^13^. Together, these contrasting transcriptional programs suggest that distinct environmental inputs contribute to phenotypic divergence between aerial and subterranean fruit.

The allo-octoploid genome of *C. chenopodiifolia* combines four sub-genomes of eight chromosomes each, creating redundancy and opportunities for duplicated genes to diversify^14,19^. Divergence of gene function following duplication is a major evolutionary force that commonly underlies trait evolution^20,21^. In *C. hirsuta* for example, leaf shape evolution involved tandem gene duplication followed by divergence in both *cis*-regulatory and protein-coding functions^22,23^. Whether, and precisely how, whole genome duplication via allopolyploidy contributed to the evolution of amphicarpy in *C. chenopodiifolia* remains unclear.

We find that light reprograms underground fruit to explode by switching SCW patterning from uniform to polar in end*b* cells, revealing remarkable developmental plasticity in *C. chenopodiifolia*. We exploit this light response to identify REVOLUTA (REV) as a transcription factor that controls end*b* cell identity, SCW deposition and organ polarity in Arabidopsis and *C. hirsuta* fruit. We further show that *REV* function has been shaped by gene and whole genome duplication in *Cardamine*, and that *REV* over-expression is sufficient to convert *C. hirsuta* end*b* SCWs from polar to uniform, transforming explosive fruit into non-explosive fruit. Together, these findings link light-dependent developmental plasticity and *REV* function to transitions in seed dispersal strategies.

## Results

### Developmental plasticity in response to light

Explosive fruit are borne on aerial shoots in the amphicarpic species *C. chenopodiifolia* and launch seeds by ultra-fast coiling of the fruit valves (Fig. 1A)^13^. Non-explosive fruit develop from subterranean flowers and the valves of these fruits do not coil (Fig. 1A)^13^. The extreme contrast between above- and below-ground environments raises the question of whether the development of explosive vs non-explosive fruit is regulated by environmental factors. We focused on the absence of light as a defining characteristic of subterranean growth. To test for developmental plasticity, we exposed subterranean fruit to light using transparent growth modules with removable light-proof covers (Fig. S1A). When subjected to light, the subterranean fruit greened and the valves explosively coiled (Fig. 1A-B). Therefore, the derived trait of non-explosive fruit in subterranean fruit switched back to explosive in response to light.

**Figure 1:**
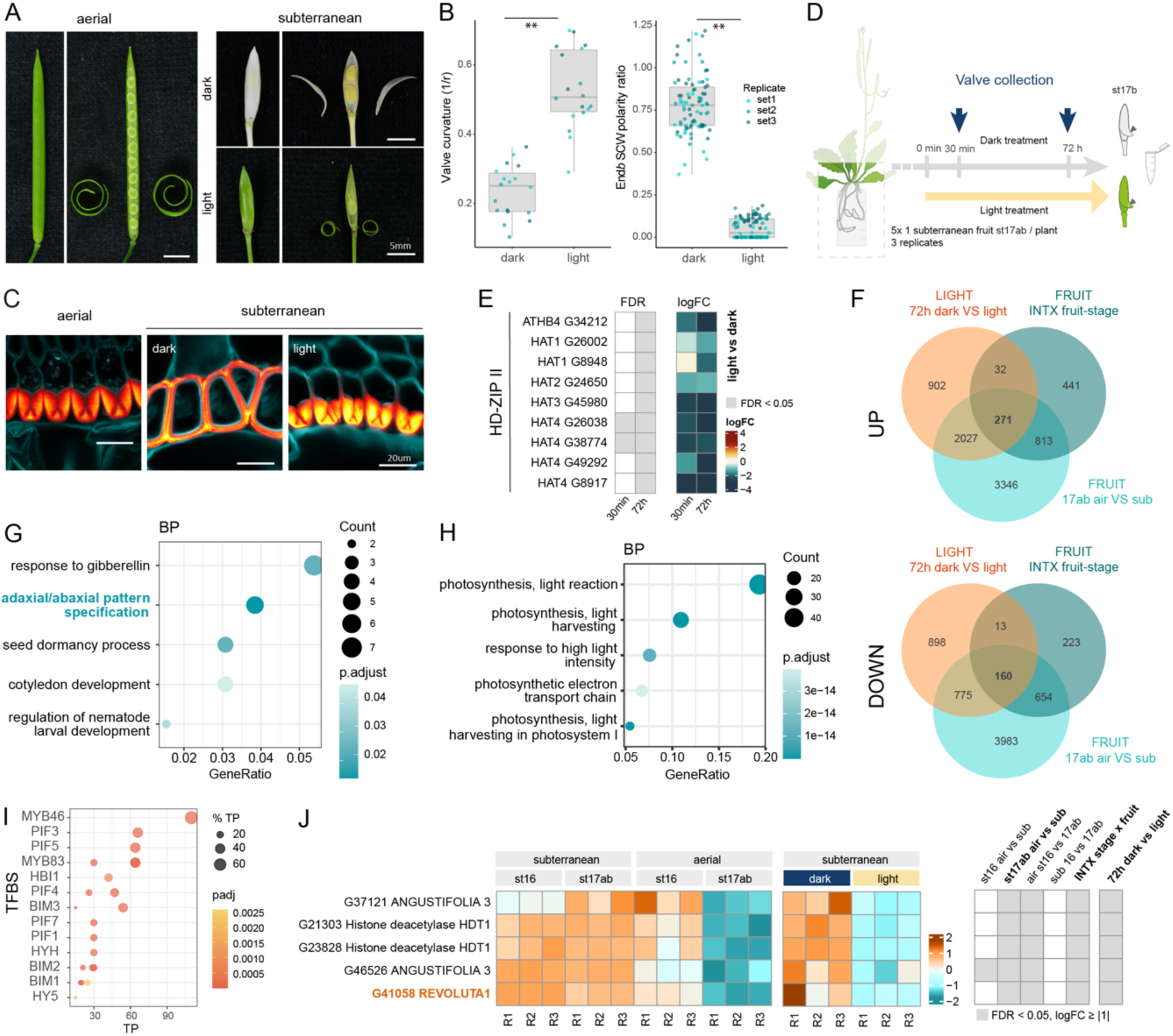
Light-responsive switch to produce explosive seed pods in *C. chenopodiifolia* **(A)** Aerial and subterranean *C. chenopodiifolia* fruit shown intact and with valves removed. Valves coil explosively in aerial fruit and subterranean fruit exposed to light. **(B)** Box plots of valve curvature (n=36) and end*b* SCW polarity ratio (n=175) in subterranean fruit grown in dark or light. Colors indicate 3 independent experiments; ** statistically significant differences at p < 0.01 using Student’s t-test; p = 1.26e-10 (valve curvature), p = 1.27e-30 (end*b* SCW polarity). **(C)** End*b* cells in transverse sections of stage 17b aerial fruit (left) and subterranean fruit grown in dark or light. Sections stained for cellulose (calcofluor white, cyan) and lignin (basic fuchsin, Red Hot LUT). **(D)** Experimental design for RNA-seq of stage 17ab subterranean fruit valves following 30 min or 72 hours of light vs dark treatment (3 replicates of 10 valves pooled from 5 different plants per sample). **(E)** *HD-ZIPII* genes down-regulated in subterranean fruit valves in response to 30 min or 72 h light vs dark shown as log_2_ fold change. **(F)** Venn diagrams of DEGs (log fold-change (logFC) ≥ |1|, false discovery rate (FDR) < 0.05) up or down-regulated between subterranean fruit valves exposed to dark vs light for 72 h (LIGHT 72h dark VS light), aerial vs subterranean fruit valves at stage 17ab (FRUIT 17ab air VS sub) and the interaction of fruit type and fruit stage (FRUIT INTX fruit-stage). **(G-H)** Selected GO terms enriched in the 160 down-regulated DEGs (G) and in the 271 up-regulated DEGs (H) in common for the comparisons depicted in (F). **(I)** Selected light-related transcription factor binding site (TFBS) motifs enriched in the 5 kb promoters of downregulated DEGs in common for comparisons between subterranean fruit valves in response to 72 h light vs dark (LIGHT 72h dark VS light), aerial vs subterranean fruit stage 17ab (FRUIT 17ab air VS sub) and interaction of fruit type and fruit stage (FRUIT INTX fruit-stage). TP: true positive sequences; padj: adjusted p-value. **(J)** Heat map of DEGs contained in the “adaxial/abaxial pattern specification” GO term highlighted in (F). Grey boxes (E, J) indicate statistically significant differences (logFC ≥ |1|, FDR < 0.05). Scale bars: 5 mm (A), 20 µm (C).

To investigate the basis for this switch to explosive valve coiling, we analyzed the pattern of lignified SCW deposition in end*b* cells. In contrast to the polar SCW pattern in end*b* cells of aerial fruit, subterranean fruit have a uniform pattern of SCW deposition (Fig. 1C, Fig. S1M-P)^13^. However, in response to light, SCW patterning switched in subterranean fruit to produce a polar SCW on the adaxial side of end*b* cells, mimicking explosive aerial fruit (Fig. 1B-C). Exposure to light also subtly shifted other subterranean fruit traits towards aerial fruit trait values. For example, fruit width and pedicel length were reduced, while seed number was increased (Fig. S1B-E). In summary, light triggered polar SCW deposition in the end*b* cell layer of non-explosive subterranean fruit valves and a switch to explosive coiling.

The plasticity of SCW patterning in response to light was surprising, since lignin is a highly stable cell wall polymer and its pattern of deposition is developmentally regulated in end*b* cells^12,13^. To investigate the lability of SCW patterning, we exposed subterranean fruit to light at stage 17ab, when uniform SCW deposition had already initiated (Fig. S1F). At time point 0, lignin deposition had initiated from cell corners, and a uniformly thickened SCW was observed after 72 hours in the dark (Fig. S1F). In contrast, after 72 hours in light, SCW thickening was confined to the adaxial side of end*b* cells with the typical hinged pattern found in aerial explosive fruit (Fig. S1F). Areas of thin lignification observed outside of this polar pattern likely represented the lignin deposited prior to light treatment. These results indicate that light can reprogram the pattern of lignin deposition during SCW formation.

We exploited this light response to identify genes controlling end*b* development and SCW patterning via RNA sequencing. Using the same fruit staging as above, we removed the light-proof covers and marked subterranean fruit at stage 17ab as time point 0, and harvested valves from these fruits after 30 min or 72 h exposure to light (Fig. 1D, Fig. S1G). In parallel, control plants were transferred to a dark room at time point 0 and valves harvested under a safe light from stage 17ab fruit after 30 min, while stage 17b fruit valves were harvested as control samples at the 72-h time point. Total RNA was extracted for Illumina sequencing and short reads were mapped to a *C. chenopodiifolia* long read reference transcriptome^13^. Very few significant differentially expressed genes (DEGs; false discovery rate of < 0.05 and minimum fold change of 2) were found after 30 min light treatment (4 DEGs) (Fig. S1H, Table S1). Two of these rapidly down-regulated genes were homeologs of the light-regulated class II homeodomain leucine zipper (HD-ZIPII) transcription factor *HAT4/ATHB2* (Fig. 1E)^24–28^. Many more DEGs were found after 72-h light treatment (5078 DEGs, 3232 up-regulated in light, 1846 down-regulated) (Fig. S1I-K) and amongst the down-regulated DEGs were nine *HD-ZIPII* genes, including four *HAT4* homeologs, two *HAT1* homeologs, *HAT2, HAT3* and *ATHB4* (Fig. 1E, Fig. S1L). Therefore, *HD-ZIPII* genes, including the direct PHYTOCHROME-INTERACTING FACTOR (PIF) target *HAT4*^29^, are rapidly repressed by light during the switch to polar SCW patterning and explosive coiling in subterranean fruit valves.

To try and increase our sensitivity to identify genes involved in the switch to polar end*b* SCW patterning, we intersected the DEGs found 72 h after light treatment with two previously published RNA-seq datasets from *C. chenopodiifolia* fruit valves^13^. We reasoned that genes involved in the SCW polarity switch may also be differentially expressed between aerial and subterranean fruit valves during end*b* lignification (FRUIT 17ab air VS sub, Fig. 1F), or in the interaction of fruit type and developmental stage (FRUIT INTX fruit-stage, Fig. 1F). Among the genes in common to the three sets of DEGs, 271 of these genes were coordinately up-regulated and 160 genes were coordinately down-regulated across all three comparisons (Fig. 1F, Table S1). Light-related transcription factor binding site motifs were enriched in the 160 down-regulated DEGs, including PIF binding motifs (Fig. 1I, Table S1). Gene Ontology (GO) processes enriched in up-regulated genes were dominated by photosynthesis, but were more diverse in down-regulated genes and included “adaxial/abaxial pattern specification” (Fig. 1G-H, Table S1).

Since light switched the SCW patterning in subterranean fruit from uniform to the adaxial side of end*b* cells, we analyzed the expression of five genes linked to the GO term “adaxial/abaxial pattern specification”, including two homeologs of histone deacetylase *HDT1*, two homeologs of *ANGUSTIFOLIA 3* and one homeolog of the class III HD-ZIP transcription factor *REVOLUTA (REV).* These genes were highly expressed in subterranean fruit valves in the dark, but strongly down-regulated in response to light (Fig. 1J). In aerial fruit valves, these genes were highly expressed prior to lignification (stage16), but strongly down-regulated during end*b* lignification (stage 17ab) (Fig.1J). Therefore, these genes were down-regulated in fruit valves that deposited a polar SCW and explosively coiled. We speculated that such genes might function in end*b* development and SCW patterning. *REV* was an attractive candidate gene since it regulates lateral organ polarity by controlling adaxial cell fate^30^. Moreover, the class II HD-ZIP genes that we identified as rapidly repressed by light, are transcriptional targets and also protein interacting partners of REV in Arabidopsis^31–33^. Therefore, we decided to investigate *REV* function in fruit valves, particularly in end*b* identity and SCW pattern formation.

### *REVOLUTA* regulates endocarp *b* identity in Arabidopsis

We first investigated *REV* function in the non-explosive fruit valves of Arabidopsis, which have uniform SCW patterning in the end*b* (Fig. 2A). An Arabidopsis *REV::REV-mVenus* functional fusion protein^34^ was nuclear-localized in the endocarp *a* and *b* cell layers of the fruit valve (Fig. 2B), as well as in vascular bundles (Fig. S2A). *REV* is a member of a small gene family of HD-ZIPIII transcription factors that are post-transcriptionally regulated by the microRNAs (miR) miR165/6 in Arabidopsis^30,35–39^. Both mature miR165 and miR166 accumulated in Arabidopsis fruit valves, with miR166 being more abundant (Fig. S2B, Table S2). The spatial expression pattern of *MIR166A::erGFP*^40^ was perfectly complementary to *REV::REV-mVenus*, marking the exocarp and mesocarp as abaxial tissues (WT, Fig. 2C, Fig. S2C). Therefore, based on the localization of *REV::REV-mVenus*, we identified that end*b* cells have adaxial identity in Arabidopsis fruit valves.

**Figure 2:**
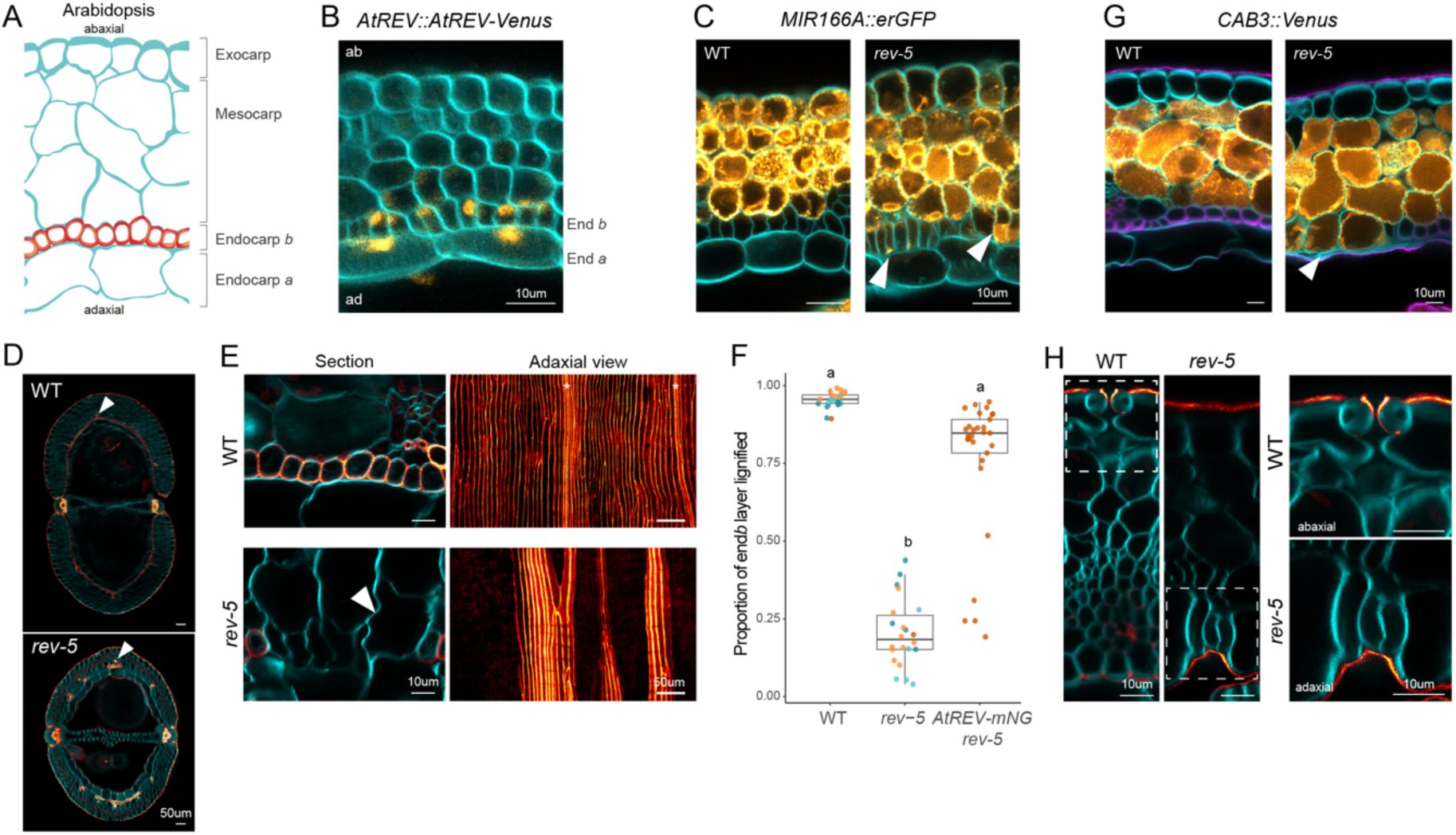
*REVOLUTA* regulates endocarp *b* cell identity in Arabidopsis **(A)** Schematic of Arabidopsis fruit valve in transverse section. **(B)** *AtREV::AtREV-mVenus* signal (Orange Hot LUT) and cellulose (cyan) in transverse section of an Arabidopsis stage 12 fruit valve (n=8). **(C)** *MIR166A::erGFP* expression (Orange Hot LUT) and cellulose (cyan) in transverse sections of Arabidopsis WT and *rev-5* stage 11 fruit valves (n=3). **(D-E)** Arabidopsis wild-type (WT) and *rev-5* mutant fruit at stage 17b stained for cellulose (cyan) and lignin (Red Hot LUT) (n = 21 WT, 22 *rev-5*). Transverse sections of whole fruit (D, arrows point out xylem vessels in representative vascular bundles) and end*b* cells (E, Section, arrow indicates mesocarp cell in end*b* cell layer in *rev-5*). Maximum projection of adaxial surface of end*b* cell layer (E, Adaxial view, stars indicate additional signal from underlying xylem vessels). **(F)** Boxplot showing ratio of the end*b* cell layer surface that is lignified in WT (n = 21), *rev-5* (n = 22) and *rev-5; AtREV::AtREV-mNG* (n = 28); colors indicate independent experiments. Different letters denote statistical significance (p = 1.61e-41) using Kruskal-Wallis test and Dunn’s multiple comparisons test (subset of analysis presented in Fig. 5I). **(G)** *CAB3::3xVenus-RCl2A* expression (Orange Hot LUT) together with cellulose (cyan) and lignin (magenta) staining in transverse sections of Arabidopsis WT and *rev-5* stage 17b fruit valves; arrow indicates mesocarp cells in end*b* cell layer in *rev-5* (n = 4 each). **(H)** Arabidopsis WT and *rev-5* stage 17b fruit valves stained for cellulose (cyan) and lignin (Red Hot LUT) showing stomata in the exocarp cell layer in WT and in the end*a* cell layer in *rev-5*; zoom-in of dashed regions shown (right). Staining: cellulose: calcofluor white (cyan); lignin: basic fuchsin (Red Hot LUT or magenta). Scale bars: 10 µm (B-C, E left panels, G-H), 50 µm (D, E right panels).

To test whether *REV* is required for end*b* development in Arabidopsis, we compared *rev-5* mutant fruit with wild type and found that lignified end*b* cells were mostly absent from mature *rev-5* fruit (Fig. 2D-E). The few lignified cells that formed in the end*b* layer of *rev-5* fruit were usually close to vascular bundles (arrow, Fig. 2D). For quantitative measurements, we stained valves for lignin and cellulose and imaged the adaxial surface of the end*b* layer to visualize the SCWs (Fig. 2E, Fig. S2E). In wild-type valves the complete end*b* surface stained, indicating that every end*b* cell had a SCW, while less than 20% of the end*b* surface stained in *rev-5* (Fig. 2F). We found the same loss of end*b* SCWs in the *rev-9* allele (Fig. S2F). We complemented the end*b* phenotype of *rev-5* by transformation with *REV::REV-mNeonGreen* (Figs. 2F) and also by crosses to *REV::REV-mVenus* (see methods). These results indicate a previously unrecognized role for *REV* in end*b* development.

To determine whether loss of end*b* cells in *rev-5* represents loss of adaxial cell identity, we analyzed cell types and marker gene expression. Non-lignified cells in the end*b* cell layer of *rev-5* valves were reminiscent of mesocarp cells (arrow, Fig. 2E; asterisks, Fig. S2G). To mark mesocarp cell identity, we used a reporter gene for the chlorophyll a/b-binding protein *CAB3* (*CAB3::3xVenus*) that expressed specifically in mesocarp cells and was excluded from endocarp, exocarp and vascular bundles in the valve (WT, Fig. 2G, Fig. S2D). In *rev-5* valves, we observed *CAB3-*expressing cells in the end*b* layer (arrow, Fig. 2G), indicating that end*b* cells had adopted an abaxial mesocarp cell fate. Expression of *MIR166A::erGFP* confirmed that end*b* cells had acquired abaxial fate in *rev-5* (Fig. 2C). Similarly, the paired guard cells that form stomata are only present in the abaxial exocarp of wild-type fruit valves, yet we found guard cells in the endocarp *a* layer on the adaxial side of *rev-5* valves (Fig. 2H, Fig. S2H-I). These findings indicate that a switch from adaxial to abaxial cell fate in endocarp layers of *rev-5* valves contributes to the loss of lignified end*b* cells.

Despite the strong effect of *REV* on end*b* development, we considered the possibility that other *HD-ZIPIII* genes might also contribute to this process. However, we did not detect any loss of lignified end*b* cells in multiple mutant combinations carrying alleles of *PHABULOSA (PHB)*, *PHAVOLUTA (PHV), CORONA (CNA)* or *ATHB8* (Fig. S2J-K). In heterozygous *phb phv rev-5* +/-segregants^41^, we observed small clusters of non-lignified end*b* cells that were not present in *phb phv* double mutants (Fig. S2J), but we were unable to further evaluate this interaction because *phb phv rev-5* triple mutants did not survive to produce fruit^30,42^. Taken together, these results indicate that *REV* regulates end*b* cell identity and SCW formation in a largely non-redundant manner in Arabidopsis.

### REVOLUTA redundancy in C. hirsuta

To investigate *REV* function in the explosive fruit of *C*. *hirsuta*, we generated fluorescent protein fusions and CRISPR-Cas9 loss-of-function mutants. *ChREV::ChREV-mNG* was nuclear-localized in endocarp *a* and *b* cells, similar to Arabidopsis, and also in an adjacent layer of small mesocarp cells that is absent in Arabidopsis (Fig. 3A-B). Interestingly, this cell layer is often lignified in subterranean fruit valves of *C. chenopodiifolia* (Fig. S1F)^13^. Therefore, based on the localization of *ChREV::ChREV-mNG*, the lignified end*b* cells in *C. hirsuta* fruit valves have adaxial identity.

**Figure 3:**
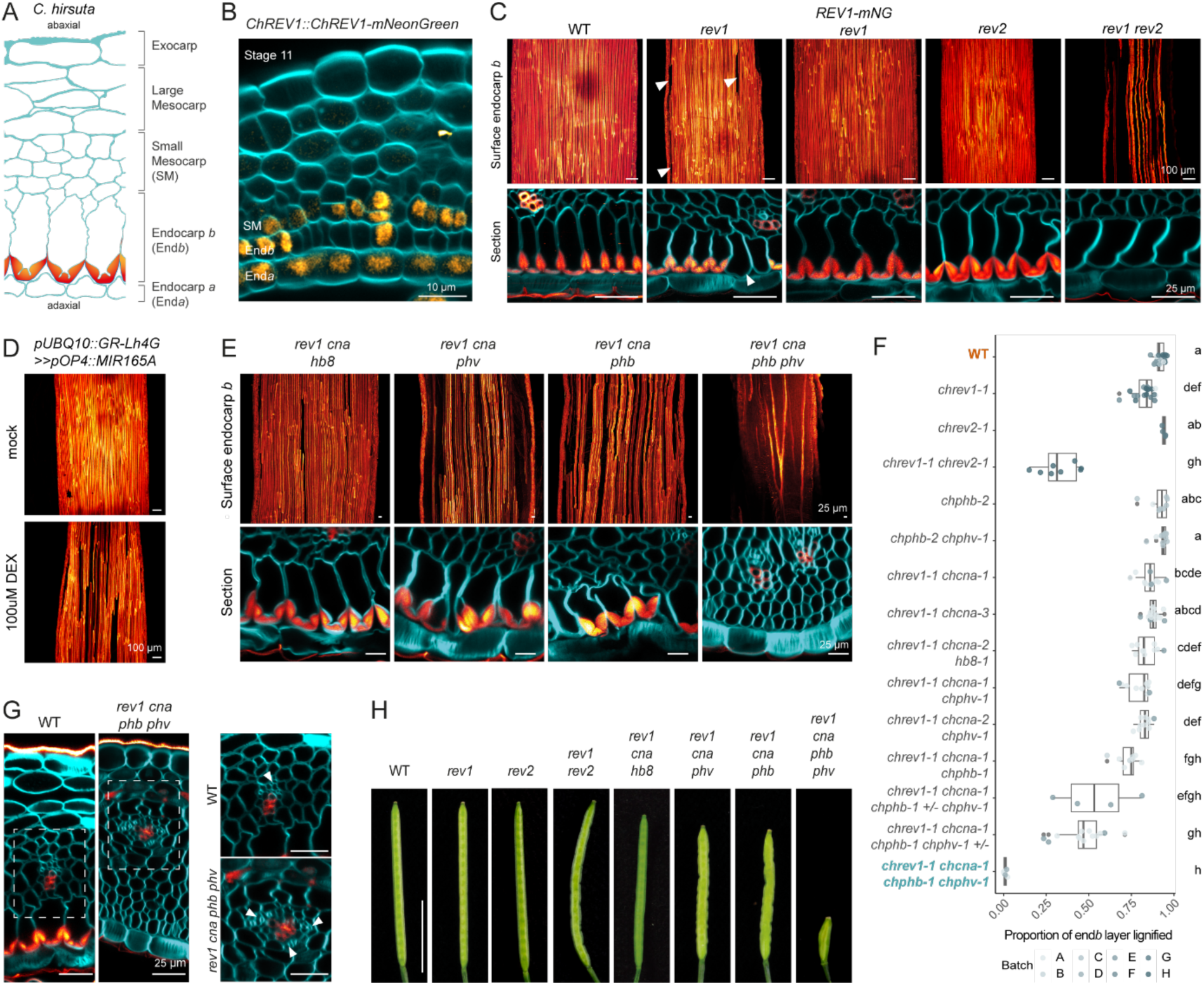
Multiple *HD-ZIPIII* genes regulate endocarp *b* cell identity in *C. hirsuta* **(A)** Schematic of *C. hirsuta* fruit valve in transverse section. **(B)** *ChREV1::ChREV1-mNeonGreen* (*mNG*) signal (Orange Hot LUT) and cellulose (calcofluor white, cyan) in transverse section of a *C. hirsuta* stage 11 fruit valve (n=5). **(C-E)** *C. hirsuta* fruit valves at stage 17b stained for cellulose (calcofluor white, cyan) and lignin (basic fuchsin, Red Hot LUT) shown as maximum projections of the adaxial surface of the end*b* cell layer or transverse sections. (C) Wild-type (WT) (n=23), *chrev1-1* (n=18) and *chrev1-1; ChREV1::ChREV1-mNeonGreen* (n=10 independent T1 lines), *chrev2-2* (n=4) and *chrev1-1 chrev2-*1 (n=8) valves; arrows indicate missing SCW. (D) *UBQ10::Gr-Lh4G>>pOP4::ChMIR165A* after mock or 100µM DEX treatment (n=27 independent T1 lines). (E) *HD-ZIPII* multiple mutants: *chrev1-1 chcna-2 chhb8-1* (n=10)*, chrev1-1 chcna-1 chphv-1* (n=10)*, chrev1-1 chcna-1 chphb-1* (n=8)*, chrev1-1 chcna-1 chphb-1 chphv-1* (n=6). **(F)** Boxplot showing ratio of the end*b* cell layer surface that is lignified in *C. hirsuta HD-ZIPIII* single and multiple mutant combinations (total n=157). Different letters denote statistical significance (p = 1.4e-19) using Kruskal-Wallis test and Dunn’s multiple comparisons test. **(G)** Vascular bundles in valve transverse sections of WT and *chrev1-1 chcna-1 chphb-1 chphv-1* quadruple mutant; zoom-in of dashed regions shown (right). **(H)** Mature fruit of *C. hirsuta* WT, *rev1-1, rev2-2, chrev1-1 chrev2-1*, *chrev1-1 chcna-2 chhb8-1, chrev1-1 chcna-1 chphv-1, chrev1-1 chcna-1 chphb-1, chrev1-1 chcna-1 chphb-1 chphv-1* (n=4-5 fruit per genotype). Scale bars: 10 µm (B), 25 µm (C, E, G), 100 µm (C, D), 1 cm (H).

We characterized the mutant phenotype of a *C. hirsuta rev-1* allele with a 10 bp deletion induced by CRISPR-Cas9, predicted to produce a truncated protein of only 54 amino acids (Fig. S3A-B). We found gaps in the end*b* layer where cells failed to form a lignified SCW (Fig. 3C). This phenotype was fully complemented by *ChREV::ChREV-mNG* expression (Fig.3C), indicating that the fusion protein is functional and the end*b* defect is caused by loss of *REV* function. The formation of lignified, interfascicular fibers was reduced in the stem of *C. hirsuta rev-1* (Fig. S4A), similar to Arabidopsis *rev* mutants^35,43^. However, loss of *REV* seems to have overall weaker effects in *C. hirsuta* compared to Arabidopsis, suggesting redundancy in gene function.

Since functional redundancy could reflect *REV* gene duplication, we searched the *C. hirsuta* genome and discovered a second *REV* gene on chromosome 5 (Fig. 5A). We called this duplicate gene *REV2*, and designated the gene orthologous to Arabidopsis *REV* as *REV1*. These *C. hirsuta* REV proteins share 96% sequence similarity and diverge in their START and MEKHLA/PAS-like domains (Fig. S3C)^44,45^. We generated six *rev2* alleles by CRISPR-Cas9 that produced either short truncated proteins or a large deletion that included the homeodomain (Fig. S3A-B). We analyzed *rev2* mutants as either single or double mutants with *rev1*. Single *rev2* mutants did not affect the fruit end*b* (Fig. 3C, F, Fig. S4B-C). Yet, *rev1 rev2* double mutants mostly lacked end*b* SCWs (Fig. 3C, Fig. S4B-C). Quantification of SCW staining in whole valves showed that only 31% of the end*b* surface stained in *rev1 rev2* double mutants (Fig. 3F). End*b* cell identity and organ polarity were unaffected in *rev1 rev2* fruit valves (Fig. 3C, S4B-C). However, these plants had reduced stature and early reproductive arrest, often producing sterile filaments in place of flowers, and leaves with fewer leaflets (Fig. S4D-F). Thus, *REV1* and *REV2* act redundantly in *C. hirsuta* to control end*b* SCW formation and other developmental processes.

Compared to *REV* function in the end*b* of Arabidopsis fruit, duplicate *REV* genes in *C. hirsuta* control SCW formation but not cell identity. This suggests that functions in cell identity and organ polarity may be redundantly shared with other *HD-ZIPIII* family members in *C. hirsuta*. Consistent with this, *REV1* and *PHB* were strongly co-expressed in *C. hirsuta* valves, but not in Arabidopsis (Fig. S5A-B, Table S2). As a functional test, ectopic expression of *C. hirsuta MIR165A* increased the amount of non-lignified gaps in the end*b* cell layer relative to *C. hirsuta rev1*, indicating that multiple miR165-silenced *HD-ZIPIII* genes contribute to end*b* development (dexamethasone-inducible *UBQ10::GR-Lh4G>>pOP4::ChMIR165A*, Fig. 3D, Fig. S5C, Table S2).

To further investigate *HD-ZIPIII* gene function, we used CRISPR-Cas9 gene editing to target *ChPHB, ChPHV, ChCNA* and *ChHB8* in the *C. hirsuta rev1-1* background, and also *ChPHB* and *ChPHV* in a wild-type background to avoid potential lethality in combination with *rev1*. We obtained mutants for combinations of all five *HD-ZIPIII* genes, targeting sequences in the first four exons and producing premature stop codons upstream of the START and the MEKHLA domains (Fig. S3A-B). The quadruple *C. hirsuta* mutant *rev1 cna phb phv* completely lacked lignified end*b* cells, demonstrating that these four *HD-ZIPIII* genes act redundantly to regulate end*b* cell identity in *C. hirsuta* (Fig. 3E-F).

We assessed organ polarity in the fruit valves produced by the *rev1 cna phb phv* quadruple mutant in *C. hirsuta*. Adaxial cell types, such as end*b*, were replaced by small mesocarp cells, and vascular bundles were radialized, with phloem surrounding xylem (Fig. 3E, G). These phenotypes are consistent with abaxialization of the fruit valve. Leaves produced by these small plants were also trumpet-shaped (Fig. S5F-G), similar to the abaxialized leaves described in allelic combinations of *rev phv* and *rev phv phb/+* in Arabidopsis^42^. Therefore, *ChREV1, ChPHB, ChPHV* and *ChCNA* regulate adaxial polarity of *C. hirsuta* fruit valves, including end*b* cell identity.

The formation of end*b* SCWs was significantly reduced when triple *rev1 cna phb* or *rev1 cna phv* mutants were additionally heterozygous for either *phv* or *phb* (Fig. 3E-F, Fig. S5D-E). Thus, when plants are wild type for *ChREV2*, a redundant role is revealed for the three *HD-ZIPIII* genes *ChPHB, ChPHV* and *ChCNA* acting with *ChREV1* to regulate end*b* SCW formation. Multi-loculate fruit are produced by these genotypes, and also by *rev1 cna phb phv* quadruple mutants, indicating that loss of these *HD-ZIPIII* genes also affects carpel development (Fig. S5E).

Complete loss of end*b* cells in *C. hirsuta rev1 cna phb phv* mutants prevented valve coiling, whereas fruit that lacked a considerable amount of the lignified end*b* layer still developed normally, but buckled along their edge at maturity (*rev1 rev2* and triple mutants of *rev1 cna phb* or *phv*, Fig. 3H, Figs. S4G, S5H). This buckling phenotype had been previously described in other *C. hirsuta* mutants with reduced lignification of the end*b* SCW and associated with reduced seed dispersal range^12^. Therefore, the regulation of end*b* identity and SCW deposition by *REV1*, acting redundantly with *REV2* and other *HD-ZIPIII* genes, is important for explosive seed dispersal in *C. hirsuta*.

### REVOLUTA is sufficient to switch endocarp *b* SCW pattern in *C. hirsuta*

We previously showed that switching the end*b* SCW pattern in subterranean fruit of *C. chenopodiifolia* from uniform to polar in response to light is sufficient to induce explosive valve coiling and is associated with *REV* down-regulation (Fig. 1A-C, J). Therefore, we hypothesized that up-regulating *REV* might be sufficient to do the opposite and switch the end*b* SCW pattern in *C. hirsuta* from polar to uniform. To test this idea, we overexpressed a miRNA-resistant version of *C. hirsuta REV1* in end*b* cells using a DEX-inducible *ChPER66::GR-Lh4G/Op4::ChREV1r-GFP* (*ChPER66::LhGR>>ChREV1r-GFP*) construct (Fig. S6A). Daily DEX treatment of stage 15 fruit in these transgenic plants during 7 days of development was sufficient to completely switch the end*b* SCW from polar to uniform (Fig. 4A-B).

**Figure 4:**
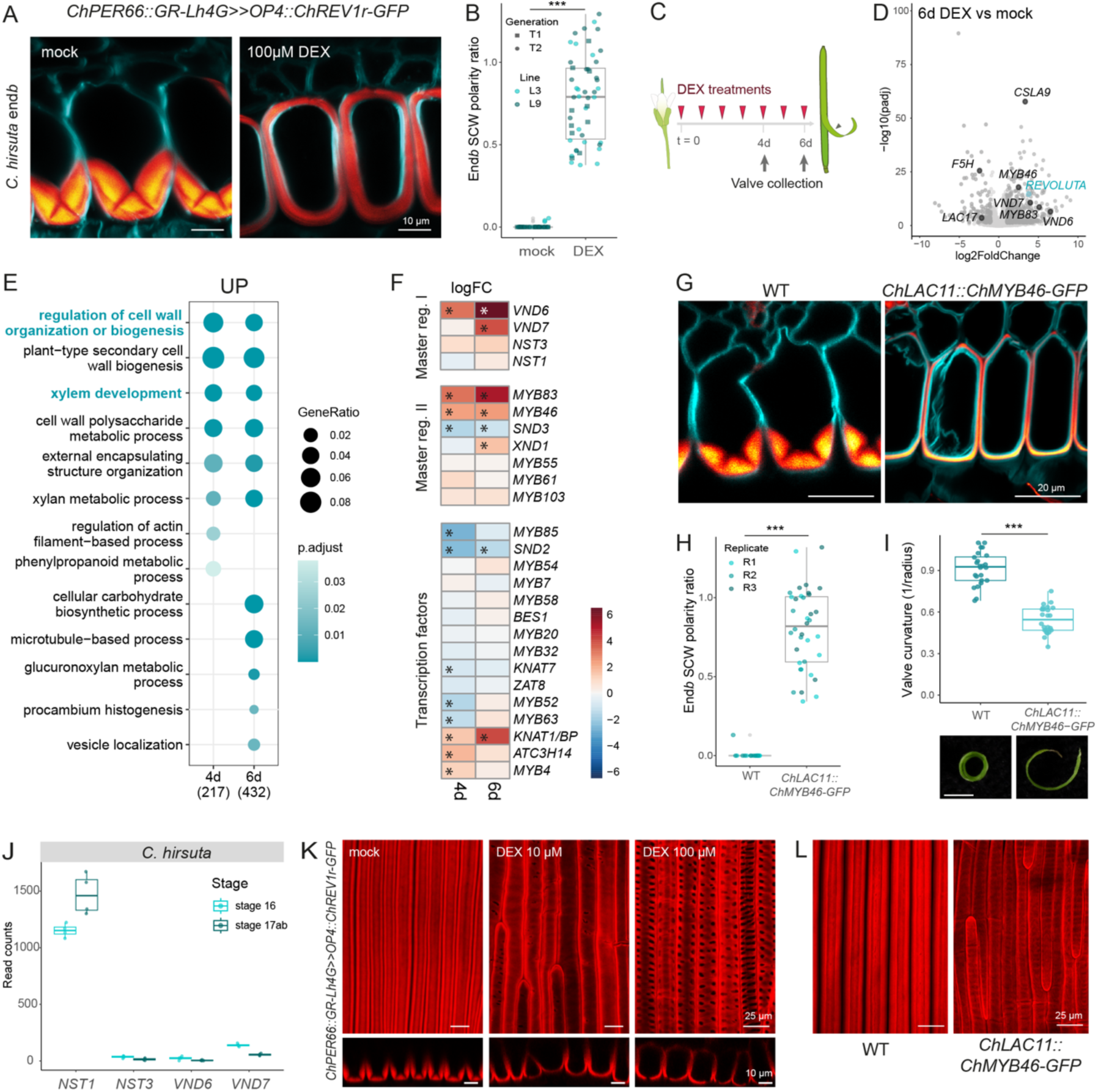
*REVOLUTA* is sufficient to switch endocarp *b* SCW pattern in *C. hirsuta* **(A)** *C. hirsuta* end*b* cells in *ChPER66::GR-Lh4G>>pOP4::ChREV1r-GFP* stage 17b fruit valves after 7-day mock or 100 µM DEX treatment stained for cellulose (calcofluor white, cyan) and lignin (basic fuchsin, Red Hot LUT) (n= 8 independent transgenic lines). **(B)** Boxplot showing end*b* SCW polarity ratio in *ChPER66::GR-Lh4G>>pOP4::ChREV1r-GFP* after 7-days treatment with mock or 100 µM DEX (total n=200, 10 cells/valve in 2 independent transgenic lines at T1 or T2 generations). Statistically significant difference (p = 9.120 e-20) using Wilcoxon Mann-Whitney test. **(C-E)** RNA-seq following DEX (100 µM) induction of *C. hirsuta ChPER66::GR-Lh4G>>pOP4::ChREV1r-GFP* fruit valves showing experimental design (C), volcano plot of DEGs (logFC) ≥│1│, padj < 0.05) at six-day (6d) time point (D) and selected GO terms enriched in up-regulated DEGs at 4d (217 genes) and 6d (432 genes) (E). **(F)** Heat map of log_2_ fold change (logFC) for master regulator genes involved in SCW formation in response to DEX induction of *C. hirsuta ChPER66::GR-Lh4G>>pOP4::ChREV1r-GFP* fruit valves; * statistically significant differences (logFC ≥ |1|, padj < 0.05). **(G)** *C. hirsuta* end*b* cells in wild-type (WT) and *ChLAC11::ChMYB46-GFP* stage 17b fruit valves stained for cellulose (calcofluor white, cyan) and lignin (basic fuchsin, Red Hot LUT) (n=3). **(H)** Boxplot showing endb SCW polarity ratio for WT and *ChLAC11::ChMYB46-GFP* fruit (total n=55). Statistically significant difference (p = 7.598 e-10) using Wilcoxon Mann-Whitney test. **(I)** Box plot of valve curvature in *ChLAC11::MYB46-GFP* and WT (n=24 valves from 12 plants per genotype). * statistically significant difference (p < 0.01) using Welch’s t-test; p = 2.983e-15. Representative coiled valve of each genotype shown below. **(J)** RNA-seq relative counts for *NST1/3* and *VND6/7* master regulators of fibers and xylem, respectively, in *C. hirsuta* fruit valves at stage 16 and 17ab. **(K)** Maximum projection of end*b* adaxial surface in *ChPER66::GR-Lh4G>>pOP4::ChREV1r-GFP* stage 17b fruit valves after treatment with mock, 10 µM and 100 µM DEX, stained for lignin (basic fuchsin, red). Optical cross-sections shown below (n=8 per treatment). **(L)** Maximum projection of end*b* adaxial surface in WT and *ChLAC11::ChMYB46-GFP* stage 17b fruit valves stained for lignin (basic fuchsin, red). Scale bars: 10 µm (A, K lower panels), 20 µm (G), 2mm (I), 25 µm (K upper panels, L).

To investigate how *ChREV1* induction affected the valve transcriptome, we harvested valves of *ChPER66::LhGR>>ChREV1r-GFP* fruit after 4 and 6 days of DEX treatment (Fig. 4C, Fig. S6B). The two time points had 560 and 668 DEGs, respectively, and *C. hirsuta REV1* was up-regulated in both time points as expected (Fig. 4D, Fig. S6C, Table S2). There were more down-regulated DEGs after 4 days and more up-regulated DEGs after 6 days with overlapping DEGs between time points (Fig. S4D). Among the up-regulated genes, GO analysis highlighted processes linked to SCW formation (Fig. 4E). Transcriptional regulation of the SCW pathway can be described as a regulatory cascade of NAC-domain and MYB-domain transcription factors that are master regulators of further transcription factors (Fig. 4F), which all together regulate structural cell wall genes (Fig. S6E)^9,10^. Master SCW regulators such as *C. hirsuta VASCULAR-RELATED NAC DOMAIN6* (*VND6*), *VND7*, *MYB46* and *MYB83* were induced in *ChPER66::LhGR>>ChREV1r-GFP* valves at both time points (Figs. 4D, F, Fig. S6C). Many genes acting downstream of these master transcription factors; for example, cellulose, hemicellulose and lignin biosynthetic genes, were differentially regulated (Fig. S6E). GO terms for lignin and hemicellulose were surprisingly enriched in down-regulated genes (Fig. S6F), possibly reflecting the thinner SCW deposited in response to *C. hirsuta REV1* compared to wild type (Fig. 4A). Overall, the largest effect of increasing *ChREV1* expression in the end*b* was to up-regulate master transcription factors that control SCW patterning and biosynthesis.

To test whether we could recapitulate the effect of *ChREV1* on end*b* SCW patterning by activating the SCW pathway, we expressed *C. hirsuta MYB46* in end*b* cells under control of the *C. hirsuta LAC11* promoter. *MYB46* and *LAC11* are both expressed in end*b* cells of *C. hirsuta* fruit (Fig. S6H)^12^, and we designed this *ChLAC11::ChMYB46-GFP* construct to create a feed-forward loop to further increase *MYB46* expression in the end*b* cell layer, based on the knowledge that *LAC11* is a direct target of MYB46 in Arabidopsis^46,47^. In *ChLAC11::ChMYB46-GFP* fruit, we observed a switch from polar to uniform deposition of the lignified SCW in end*b* cells (Fig. 4G-H) and a reduction in explosive valve coiling (Fig. 4I, Fig. S6G). Therefore, increased expression of either *REV* or *MYB46* in *C. hirsuta* end*b* cells over-rides the endogenous polar SCW patterning process, resulting in a uniform SCW and non-explosive seed dispersal.

We noticed that the GO term ‘xylem development’ was enriched in the DEGs up-regulated by DEX induction in *ChPER66::LhGR>>ChREV1r-GFP* fruit valves (Fig. 4E). Two highly up-regulated genes— *VND6* and *VND7*—are master regulators that are sufficient for metaxylem and protoxylem vessel formation^48^. These genes have low expression levels in wild-type fruit valves (Fig. 4J, Table S2). *NST1* is the master SCW regulator that is usually expressed at high levels in wild-type fruit valves of *C. hirsuta* (Fig. 4J) together with *NST3* in Arabidopsis (Fig. S6I, Table S2). Thus, *ChREV1* induction appears to reprogram the SCW transcriptome towards xylem development in *C. hirsuta* valves by activation of *VND6/7*. For example, transcription factors involved in processes specific to xylem, but not end*b* cells, such as programmed cell death, were up-regulated by *C. hirsuta REV1* (Fig. S6E).

To analyze the consequences of this transcriptional reprogramming on end*b* SCW patterning, we imaged the adaxial surface of the end*b* layer in DEX-treated *ChPER66::LhGR>>ChREV1r-GFP* fruit (Fig. 4K). Strikingly, the end*b* SCW was punctuated by holes similar to metaxylem pits (Fig.4K). Since high levels of *ChREV1*, but not *ChMYB46* (Fig. 4L), caused metaxylem patterning of the end*b* SCW, this is likely a consequence of *VND6/7* activation by ChREV1. By comparing treatments with increasing concentrations of DEX, we found a dose-dependence of *ChREV1* induction on the density and patterning of SCW pits (Fig.4K). Low levels of *ChREV1* induced few, random pits and loss of the characteristic hinged SCW pattern (10 µM DEX, Fig. 4K). Higher *ChREV1* levels triggered uniform SCW deposition and regular formation of pits aligned along the cell edges where hinges would normally form (100 µM DEX, Fig. 4K). Despite the uniform deposition of SCW, pits mostly formed on the adaxial wall of end*b* cells (Fig. S6K). These findings suggest that the SCW depletion required to form both pits and hinges might share some common patterning information. In Arabidopsis, *REV* overexpression in the *rev-10d* mutant did not alter the uniform SCW found in wild-type end*b* cells, but occasionally caused ectopic SCW deposition in a pitted pattern in endocarp *a* cells (Fig. S6J). In summary, *ChREV1* levels determine the patterning of SCW deposition in *C. hirsuta* end*b* cells.

### *REVOLUTA* gene duplication and divergence in *C. chenopodiifolia*

Our results reveal a genetic route for converting explosive to non-explosive fruit via upregulation of *REV* and its target genes, raising the possibility that *REV* may contribute to the evolution of non-explosive subterranean fruit in *C. chenopodiifolia*. To start to investigate this, we analyzed the octoploid genome of *C. chenopodiifolia* and identified eight *REV* genes, organized into two synteny groups per sub-genome (Fig. 5A). One group of four homeologs are syntenic with *REV1* located on chromosome 8 in *C. hirsuta*, while the other group is syntenic with *C. hirsuta REV2* on chromosome 5 (Fig. 5A). In sub-genome C of *C. chenopodiifolia*, *CcREV1-C* is a likely pseudo-gene, truncated in the middle of the START domain and therefore lacking both bZIP and homeobox domains (Fig. 5B). In addition, *CcREV1-C* transcripts were barely detectable throughout our RNA-seq experiments (Fig. S7A). In sub-genome B, *CcREV2-B* terminates prematurely in the middle of the START domain and lacks the terminal MEKHLA domain (Fig. 5B). All *C. chenopodiifolia REV* genes show conservation of the miR165/6 complementary site located in the START domain (Fig. 5B)^30^. In sub-genome A, *CcREV1-A* contains a T to C transition immediately 5’ to the miR165/6 complementary site, which is not likely to affect post-transcriptional regulation (Fig. 5B)^49^. Therefore, the polyploid genome of *C. chenopodiifolia* contains eight *REV* genes that fall in two syntenic groups with sequence divergence, including pseudogenization, observed across different sub-genomes.

**Figure 5:**
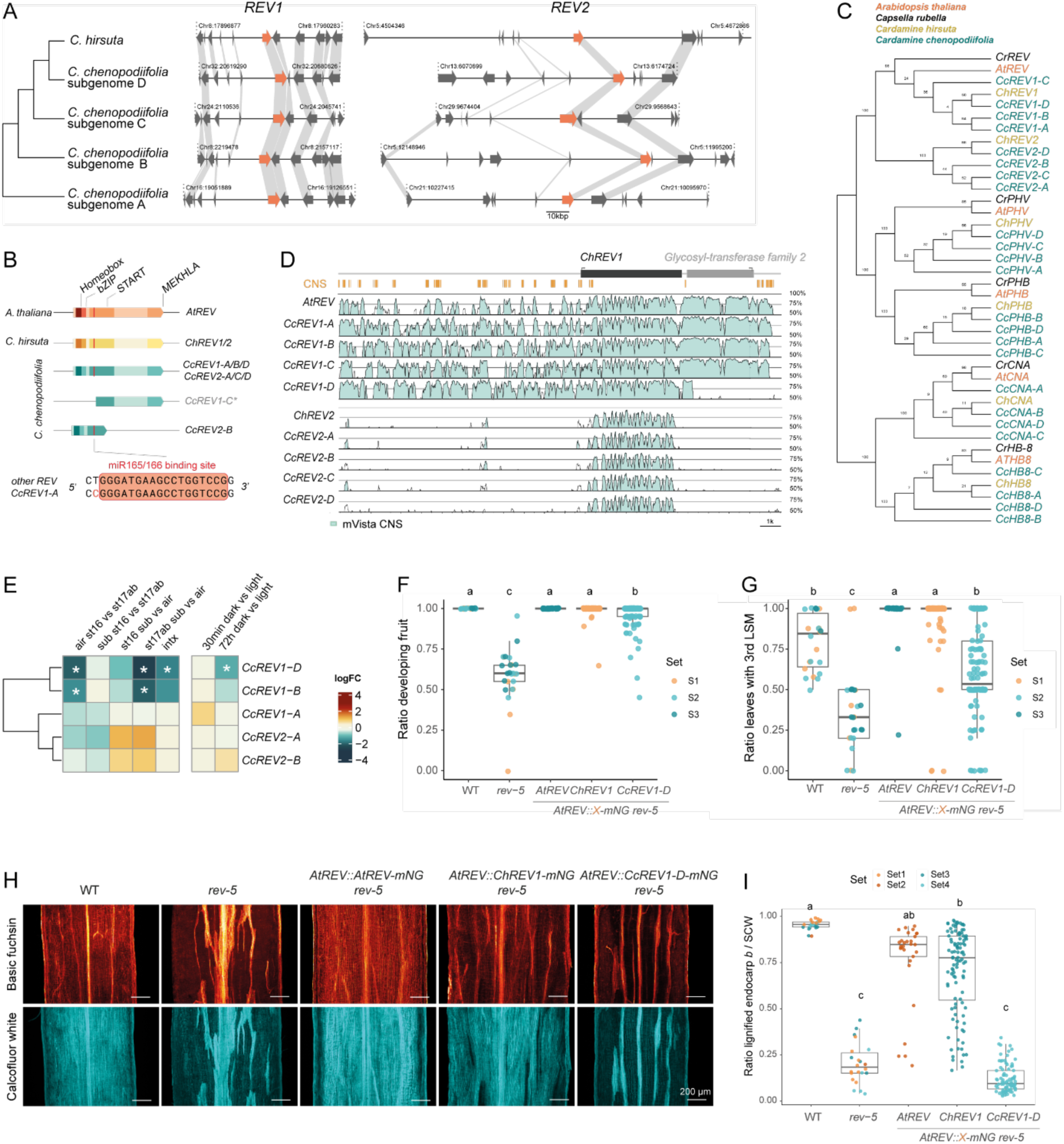
*REVOLUTA* gene duplication and divergence in *C. chenopodiifolia* **(A)** Synteny plots of *REV1* and *REV2* intervals in the diploid *C. hirsuta* genome and the four sub-genomes of *C. chenopodiifolia*. Arrows, genes; grey shading, synteny; genome coordinates shown; phylogeny as previously described^14^. **(B)** Schematic of REV proteins with conserved domains annotated in different color shades in Arabidopsis (orange), *C. hirsuta* (yellow) and *C. chenopodiifolia* (cyan); miR165/6 complementary site (red bar); *CcREV1-C* likely pseudogene (*). **(C)** Maximum likelihood cladogram of *HD-ZIPIII* gene family members in Arabidopsis (*At*), *C. hirsuta* (*Ch*), *C. chenopodiifolia* (*Cc*) and *Capsella rubella* (*Cr*). JTT model with rate heterogeneity was used and the tree with the highest log likelihood (−10552.18) shown. Numbers at each node are bootstrap values. **(D)** Conservation analysis of a 20 kb window surrounding *C. hirsuta REV1* showing CNS blocks (orange boxes) and mVISTA plots (light blue) of *ChREV1* orthologs in Arabidopsis (*AtREV*) and *C. chenopodiifolia* (*CcREV1-A/B/C/D*) and the paralogous genes in *C. hirsuta* (*ChREV2*) and *C. chenopodiifolia* (*CcREV2-A/B/C/D*). **(E)** Heat map of log_2_ fold change (logFC) values for six *C. chenopodiifolia REV* genes (*CcREV1-A/B/D* and *CcREV2-A/B*) in comparisons between aerial (air) and subterranean (sub) fruit valves at stages 16 and 17ab and the interaction of fruit type and fruit stage (INTX) *(first panel)* and subterranean fruit valves exposed to dark vs light for 30 min and 72 h *(second panel)*; * statistically significant differences (logFC ≥│1│, FDR < 0.05). **(F-I)** Transgenic complementation assays in Arabidopsis wild-type WT (n=22), *rev-5* (n=26), and *rev-5* complemented with *AtREV::AtREV-mNG* (T1, n=32)*, AtREV::ChREV1-mNG* (T2, n=135) and *AtREV::CcREV1-D-mNG* (T2, n=122). Results pooled from independent sets of experiments (S1-4). Different letters denote statistical significance at p = 1.61e-41 (F), p = 1.06e-35 (G), p = 2.79e-36 (H) using Kruskal-Wallis test and Dunn’s multiple comparisons test. **(F-G)** Ratio of successful meristem development of flowers (F, visible fruit) and axillary shoots (G, visible 3^rd^ lateral shoot meristem in secondary cauline leaves). **(H)** Maximum projection of adaxial end*b* surface showing cellulose (calcofluor white, cyan) and lignin (basic fuchsin, Red Hot LUT). **(I)** Boxplot showing ratio of the end*b* cell layer surface that is lignified in (H). Scale bars: 10 kbp (A), 200 µm (H).

A cladogram of all *HD-ZIPIII* genes in *C. chenopodiifolia* showed that *REV* is the only gene family member to have undergone additional duplication; all other genes have four homeologs (Fig. 5C). Since Arabidopsis and *Capsella rubella* contain only one *REV* gene that is orthologous to *REV1* genes in *C. hirsuta* and *C. chenopodiifolia*, it is likely that *REV2* resulted from a duplication of *REV1* that occurred in the *Cardamine* lineage (Fig. 5C). To further characterize *C. chenopodiifolia REV* genes we analyzed non-coding sequence conservation. First, we identified conserved non-coding sequences (CNS) for *REV1* in *C. hirsuta* using the Conservatory algorithm^50^. We found 109 CNS, including 44 deeply conserved CNS, which suggested that gene regulatory programs for *REV1* have been conserved over large evolutionary timescales (Fig. 5D). These CNS were embedded in blocks of sequence similarity defined by mVISTA alignments in Arabidopsis *REV* and *C. chenopodiifolia REV1* genes (Fig. 5D). A conspicuous 5.5 kb deletion immediately downstream of *CcREV1-D* removed sequences similar to 18 of the *C. hirsuta REV1* CNS (Fig. 5C). Apart from this, non-coding sequences between *REV1* genes were broadly conserved. By contrast, the lack of similarity between *REV1* and *REV2* non-coding sequences was striking (Fig. 5D). Conservatory identified only 13 CNS for *REV2* in *C. hirsuta* and mVISTA alignments showed only two peaks of similarity between *C. hirsuta REV1* and any of the *REV2* genes in *C. hirsuta* or *C. chenopodiifolia* (Fig. 5D), suggesting rapid divergence following gene duplication.

To compare expression divergence amongst duplicated *REV* genes in *C. chenopodiifolia*, we examined six genes that were present in our long read reference transcriptome (Fig. S7A). Among these genes, only *CcREV1-D* and *CcREV1-B* were highly expressed in subterranean fruit valves and differentially expressed in fruit valve datasets (Fig. 5E, Fig. S7A). Both genes were highly expressed throughout the development of dark-grown subterranean fruit and *CcREV1-D* was significantly down-regulated in response to light (Fig. 5E, Fig. S7A). In aerial fruit, both genes were significantly down-regulated at later stages of fruit valve development (Fig. 5E, Fig. S7A), similar to the pattern of *REV* gene expression in Arabidopsis and *REV1* in *C. hirsuta* (Fig. S7B-C). Therefore, expression divergence of *CcREV1-D* and *CcREV1-B* was associated with the reorganization of regulatory networks in subterranean fruit valves of *C. chenopodiifolia*.

To investigate functional divergence of the *CcREV1-D* gene, we used transgenic complementation assays. To assess protein coding function, we introduced the genomic sequences of *C. chenopodiifolia CcREV1-D*, *C. hirsuta REV1* and Arabidopsis *REV,* each driven by the Arabidopsis *REV* promoter, in the *rev-5* mutant of Arabidopsis. We compared the ability of each protein-coding sequence to complement loss of *REV* function in axillary and flower meristem formation, and lignified end*b* formation in fruit valves (Fig. 5F-I, Fig. S7D). The *REV* genes from Arabidopsis and *C. hirsuta* complemented all phenotypes, showing significantly different phenotype values to *rev-5* (Fig. 5F-I). Similar complementation of *rev-5* axillary and floral meristem phenotypes was observed by *C. chenopodiifolia CcREV1-D* (Fig. 5F-G). However, *CcREV1-D* failed to complement the formation of lignified end*b* cells in *rev-5* fruit valves (Fig. 5H-I). Therefore, *CcREV1-D* has both conserved and diverged functions. The diverged function of *CcREV1-D* in end*b* development and SCW formation is likely to reflect derived aspects of non-explosive seed dispersal in subterranean fruit of *C. chenopodiifolia*, rather than simply phylogenetic distance, given the complementation of Arabidopsis *rev-5* by *C. hirsuta REV1* (Fig. 5H-I). Taken together, *REV* duplication and functional divergence are linked to amphicarpy in *C. chenopodiifolia*.

## Discussion

Here we discovered that light can trigger underground seed pods to explode. We traced the mechanism of explosion to a light-induced switch in SCW geometry in the fruit end*b* cell layer. We identified the transcription factor REVOLUTA (REV) as necessary for end*b* cell fate and SCW formation, and sufficient to switch SCW patterning. We found that gene duplication and allopolyploidy contributed to *REV* redundancy and functional divergence between closely related species. Thus, by exploiting an understudied species with a novel seed dispersal trait, we made the advance of linking trait evolution to environment and *REV* gene function.

HD-ZIPIII transcription factors are part of a patterning mechanism that governs adaxial/abaxial polarity and radial patterning throughout plant development^30,36,42,51,52^. Fruit valves are lateral organs, homologous to leaves^53^, with distinct adaxial/abaxial polarity marked by REV expression in adaxial endocarp tissues, and *MIR166A* expression in abaxial tissues. Development of lignified end*b* cells in this adaxial domain requires *REV* in Arabidopsis. Although small clusters of end*b* cells still formed in Arabidopsis *rev-5* mutants, the lethality of *rev phb phv* mutants^30,42^ precluded further assessment of these genes’ contribution to the *rev* fruit phenotype. In contrast, *REV* gene duplication in *C. hirsuta* created redundancy and allowed the recovery of fertile *rev1 phb phv cna* quadruple mutants, which produced fruit that lacked end*b* cell identity entirely. The *rev1 rev2* double mutant in *C. hirsuta* lacked end*b* SCWs without affecting cell fate, thus separating the shared functions of Arabidopsis *REV.* More generally, recovering higher-order *HD-ZIPIII* mutants in *C. hirsuta* opens new opportunities to dissect *HD-ZIPIII* functions in plant development.

The radialized vascular bundles of *C. hirsuta rev1 phb phv cna* quadruple mutant fruit valves exhibit abaxialization, resembling the radialized cotyledons of Arabidopsis *phb phv rev* seedlings and contrasting with the adaxialized vascular bundles in stems of *rev-10d* gain-of-function mutants^30^. These findings underscore the conserved role of *HD-ZIPIII* genes in establishing radial tissue patterns across diverse plant organs, including roots, where they redundantly control xylem formation^51,54^. Endodermally produced miR165/6 degrades *PHB* mRNA at the periphery of the vascular cylinder, creating a radial PHB gradient that specifies metaxylem with pitted SCWs at the center and protoxylem at the periphery^51,55^. Similarly, high levels of *REV1* overexpression in *C. hirsuta* end*b* cells induced pitted SCWs, likely as a consequence of *VND6/7* activation^48^. Overexpression of *REV1*, or its target *MYB46*, converted the polar end*b* SCWs in *C. hirsuta* to a uniform pattern. This loss of polar SCW patterning produced non-explosive fruit, revealing a genetic mechanism that can switch seed dispersal from explosive to non-explosive.

SCW deposition in *C. hirsuta* end *b* cells was sensitive to both loss and gain of *REV* function. In *rev1 rev2* mutants, end*b* SCWs were largely absent, consistent with the established role of *REV* in the Arabidopsis SCW gene regulatory network^10^. Conversely, gain of *REV1* function in *C. hirsuta* end*b* cells activated key regulators in this network, including *MYB46* and *MYB83*, shifting SCW deposition from polar to uniform. In *C. chenopodiifolia*, light exposure induced the opposite switch—from uniform to polar SCW deposition—converting subterranean fruits from non-explosive to explosive. Notably, *REV* was among the down-regulated genes in these explosive fruits, with MYB46 and MYB83 binding sites enriched in this gene set (Fig. 1I). Together, these complementary patterns suggest that *REV* acts as a central hub in the SCW network, whose activity can be tuned in opposite directions to control end*b* SCW patterning in *Cardamine*, thereby driving transitions between explosive and non-explosive seed dispersal strategies.

Underground fruit are an evolutionary novelty of *C. chenopodiifolia* and introduce dark-activated signaling into fruit development. PHYTOCHROME-INTERACTING FACTOR (PIF) bHLH transcription factors regulate numerous target genes in germinating Arabidopsis seedlings to repress photomorphogenesis in the dark and instead promote skotomorphogenic growth through the soil^56^. PIFs are rapidly degraded within minutes upon light exposure through the binding of photoactivated phytochrome proteins^57,58^. In this way, direct PIF targets, such as the *HD-ZIPII* gene *HAT4/ATHB2*, are activated in the dark and rapidly repressed in light^29^. In subterranean fruit of *C. chenopodiifolia*, two *HAT4/ATHB2* homeologs were rapidly repressed within 30 min of light exposure and seven additional *HD-ZIPII* genes after 72 h (Fig. 1E). Moreover, PIF transcription factor binding sites were enriched in genes down-regulated in fruit valves that became explosive in response to light or differed by fruit type (Fig. 1I). Therefore, canonical light-dark signaling pathways may have been co-opted during the evolution of subterranean fruit in *C. chenopodiifolia*, enabling fruit to sense their burial in the soil and adaptively switch seed dispersal strategy in response to local soil conditions.

Interactions between REV and PIF-dependent HD-ZIPII transcription factors may link light-dark signaling to SCW patterning in subterranean fruit. In Arabidopsis, REV and HD-ZIPIIs form a regulatory loop in which REV activates *HD-ZIPII* transcription, HD-ZIPII proteins interact with REV to repress *MIR165/6* expression, and miR165/6 post-transcriptionally repress *REV*^30–33,49^. Applied to subterranean fruit, this circuitry could help maintain *CcREV1-D* expression in the dark, while light rapidly represses *HD-ZIPII* genes and shifts the balance of this network. The recent establishment of genetic transformation in *C. chenopodiifolia*^13^ now makes it possible to test these regulatory interactions using genome editing, offering a powerful route to determine how environmental cues were wired into fruit developmental networks during polyploid evolution.

Both gene duplication and allopolyploidy have driven functional diversification of *REV* in *Cardamine*. In *C. hirsuta*, *REV* paralogs have specialized to uncouple roles in end*b* SCW formation from cell identity and organ polarity, whereas in *C. chenopodiifolia*, extensive divergence among *REV* homeologs has accompanied the evolution of light-responsive seed dispersal strategies in subterranean fruit. *REV* genes have also duplicated in major cereal crops – including rice, maize and wheat – and miR165/6-resistant alleles of different paralogs in these species generate agronomically important phenotypes, such as increased grain yield and protein content^59–61^. These examples illustrate how expansion of *REV* genes through gene and genome duplication provides a rich source of genetic variation for both evolutionary innovation and crop improvement. Extending comparative studies beyond established models into polyploid species such as *C. chenopodiifolia* thus offers exciting opportunities to uncover how duplicated developmental regulators acquire new functions and generate trait diversity.

## Resource availability

### Lead contact

Requests for further information and resources should be directed to and will be fulfilled by the lead contact, Angela Hay.

### Material availability

All plasmids and plant lines generated in this study are available from the lead contact without restriction.

### Data and code availability

- Short-sequence read data and PacBio long-sequence read data for this study have been deposited in the European Nucleotide Archive (ENA) at the European Molecular Biology Laboratory’s European Bioinformatics Institute (EMBL-EBI) under accession no.: PRJEB102376, PRJEB69676.
- This paper does not report original code.
- Any additional information required to reanalyze the data reported in this paper is available from the lead contact upon request.

## Acknowledgements

We thank M. Tsiantis for comments, and together with R. Dello Ioio, C. Whitewoods, M. Heisler, M. Timmermans and N. Geldner for sharing materials, W. Faigl for technical assistance, the Max Planck-Genome-Centre Cologne for sequencing and N. Bologna for help with miRNA analysis. This work was supported by Swiss National Science Foundation fellowships P500PB_203021 and P5R5PB_225481 to A.E. and core funding of University of Bern to M.R. A.H. acknowledges support from the Max Planck Society and the Department of Comparative Development and Genetics.

## Author contributions

Conceptualization, A.E. and A.H.; Investigation, A.E., I.S., C.K., N.T. and L.B.; Bioinformatics, S.D, A.E.; Cell wall analysis, M.P.; Writing, A.E. and A.H.; Funding Acquisition, A.E., M.R. and A.H.; Supervision, A.H.

## Supplemental figures

Figure S1: Light-responsive switch to produce explosive seed pods in *C. chenopodiifolia*, related to Figure 1.

Figure S2: *REVOLUTA* regulates endocarp *b* identity in Arabidopsis, related to Figure 2.

Figure S3: Characterization of *C. hirsuta* CRISPR/Cas9 *HD-ZIPIII* alleles, related to Figure 3.

Figure S4: Phenotypic analysis of *C. hirsuta rev1, rev2* single and double mutants, related to Figure 3.

Figure S5: Phenotypic analysis of multiple *HD-ZIPIII* mutants in *C. hirsuta,* related to Figure 3.

Figure S6: *REVOLUTA* is sufficient to switch endocarp *b* SCW pattern in *C. hirsuta,* related to Figure 4.

Figure S7: *REVOLUTA* gene duplication and divergence in *C. chenopodiifolia,* related to Figure 5.

## Supplemental information

Table S1: DE analysis of *C. chenopodiifolia* data, related to Figure 1.

Table S2: DE analysis of *C. hirsuta* and Arabidopsis data, related to Figure 4.

Table S3: Primers and small guide RNA, related to STAR Methods.

## STAR Methods

### EXPERIMENTAL MODEL AND STUDY PARTICIPANT DETAILS

#### Plant material and growth conditions

*Arabidopsis thaliana* Columbia (Col-0), *Cardamine hirsuta* Oxford (Ox) herbarium specimen voucher Hay 1 (OXF)^62^ and *Cardamine chenopodiifolia* Pers. (Ipen: XX-0-MJG-19—35600)^13^ were used as wild types and the backgrounds for all genotypes. Arabidopsis mutant alleles *rev-5*, *rev-9, rev-10D, cna-2, athb8-1*, *phb-13*, *phv-11*, *phb* (SALK032108.50.25.x), *phv* (SAIL_899_C08)^63^ were previously described^30,41–43^. Arabidopsis transgenic lines *MIR166A::erGFP*^40^ and *REV::REV-VENUS*^34^ were previously described and *CAB3::3xVenus-RCl2A* (*Cre-U5-Cre_CAB3::lox-CyPET-lox-3xVenus-RCl2A,* note: this line also contains *ATML1::lox2272-CyPET-lox-tbRED-RCl2A*) was a gift from C. Whitewoods. These reporters were crossed to *rev-5* to obtain *CAB3::3xVenus-RCl2A rev-5*, *REV::REV-VENUS rev-5* and *MIR166A::erGFP rev-5* lines.

Aerial seeds were surface sterilized in 70% ethanol, 0.1% Triton-X-100 for 5–20 min, then washed in 100% ethanol and dried in sterile conditions. Seeds were plated on ½ Murashige and Skoog (MS), 500 mg l^−1^ MES and 0.8% agar medium and stratified in the dark at 4°C for 2 days (Arabidopsis) or 7 days (*C. hirsuta, C. chenopodiifolia*). Seedlings were grown for 7 days in long day conditions (16h:8h, light:dark, 20°C:18°C, fluence rate 115.9 µmol m^-^² s^-1^) in a culture room, then transferred to soil. *C. chenopodiifolia* was grown in walk-in chambers (Cologne) in long-day conditions (16h:8h, light:dark, 20°C:18°C, 65% humidity, fluence rate 85.98 µmol m^-^² s^-1^). *C. hirsuta* and Arabidopsis were grown in greenhouses (Cologne) or a culture room (Bern) in long-day conditions (16h:8h, light:dark, 20°C:18°C) or in the same walk-in chambers as *C. chenopodiifolia* (Cologne) for all RNA-seq experiments.

## METHOD DETAILS

### Plasmid construction and plant transformation

Plasmids were constructed using In-Fusion Advantage PCR cloning (Clontech), Gateway cloning (Invitrogen, Thermo Fisher Scientific, USA) and Multisite GreenGate cloning^64^. All plasmids were transformed by heat shock into *Agrobacterium tumefaciens* GV3101 strain and transformed into plants via floral dip^65^.

*AtREV::AtREV-mNG* (*mNeonGreen*)*, ChREV1::ChREV1-mNG, AtREV::ChREV1::mNG, AtREV::CcREV1-D-mNG, ChPER66::GR-LhG4>>pOP4::ChREV1r-GFP, UBQ10::GR-LhG4>>OP4::ChMIR165A, ChLAC11::ChMYB46-GFP* and *ChMYB46::NLS-3xGFP* were generated by GreenGate cloning after all BsaI sites were mutagenized as described (see Table S3 for primers)^64^. The following promoters were PCR-amplified (base pairs [bp] before ATG) in this study and inserted in a pGGA empty vector: *pAtREV* (3774bp), *pChREV1* (4744bp), *pChMYB46* (3156bp). *AtREV, ChREV1, CcREV1-D, ChMIR165A* full genomic sequences and *ChMYB46* coding sequence were amplified and cloned into pGGC entry clones. *ChREV1r* was generated by amplification of the *ChREV1* genomic sequence with a targeted mutation in the miRNA recognition site (TTTGGT**G**TCAAG instead of TTTGGT**C**TCAAG). Wild-type *ChREV1* sequence was obtained from point-targeted mutation of *ChREVr-pGGC* entry clone by In-Fusion cloning. The sequence *linker-mNeonGreen* was cloned in the pGGD empty module^11^. The promoter entry clones *pGGA-pChPER66*, *pGGA-pChLAC11* and *pGGA-pUBQ10* were previously described^12^. All entry vector combinations were cloned into the pGGZwf01 binary vector^12^. The constructs *ChLAC11::ChMYB46c-GFP*, *ChMYB46::NLS-3xGFP* and *ChREV1::ChREV1-mNG* were transformed into *C. hirsuta* wild type and *ChREV1::ChREV1-mNG* was transformed into *C. hirsuta rev1-1*. The constructs *AtREV::AtREV-mNG*, *AtREV::ChREV1-mNG* and *AtREV::CcREV1-D-mNG* were transformed into Arabidopsis *rev-5*.

*ChPER66::GR-Lh4G>>OP4::ChREV1r-GFP* and *UBQ10::GR-Lh4G>>OP4::ChMIR165A* were generated as multiple expression GreenGate cassettes using the entry vectors *pOP4-pGGA* and *GR-LhG4-pGGC*^66^ (Table S3). Entry vectors were subsequently cloned in pGGZwf01 binary vector and transformed into *C. hirsuta*.

### CRISPR-Cas9 genome editing

CRISPR/Cas9 genome editing of *C. hirsuta REV1* was performed according to^67^. A plasmid containing sgRNAs driven by AtU6 promoters in a pUC57 backbone was synthesized by GenScript Biotech (Leiden, Netherlands) and recombined by MultiSite Gateway Cloning with the destination vector pDeCas9 containing the SpCas9 sequence driven by the pPcUbi4-2 ubiquitous promoter and a Basta resistance marker^68^. This construct was transformed into *C. hirsuta* wild type to generate the *chrev1-1* allele.

A multiplex sgRNA cloning strategy was used to edit *ChPHB* and *ChPHV*, or *ChREV2* alone, in *C. hirsuta* wild type, and to edit *ChPHB, ChPHV, ChCNA* and *ChHB8,* or *ChREV2* alone, in *chrev1-1*, according to^69^. SpCas9-compatible sgRNAs were designed using CCTop (https://cctop.cos.uni-heidelberg.de^70^), CRISPRater^71^ and Geneious Prime (http://www.geneious.com/; Version 10.2.6; Biomatters Ltd., Auckland, New Zealand) (Table S3), synthesized and inserted in pRU41-pRU48 vectors containing either the pAtU6 or pAtU3 promoters. For targeting *ChPHB* and *ChPHV* (two sgRNA each) or *ChREV2* (four sgRNAs), the four corresponding entry vectors were combined by Greengate cloning into pSF464. For targeting *ChPHB, ChPHV, ChCNA* and *ChHB8* together, all entry clones were combined into pRU325. The two binary vectors were assembled by Gateway cloning into pRU295 and pRU294, which contain the *pEC1.2::zCas9i* cassette with respectively FastGreen and FastRed selection markers. The final constructs targeting *ChPHB* and *ChPHV,* or *ChREV2* alone, were transformed into *C. hirsuta* wild type, while the constructs targeting *ChPHB, ChPHV, ChCNA* and *ChHB8* together, or *ChREV2* alone, were transformed into *chrev1-1*.

Selection of CRISPR mutants was performed in T1 plants by PCR amplification and sequencing of the targeted sequences (see primers in Table S3). Sanger sequencing data was analyzed using the ICE CRISPR Analysis Tool (https://www.editco.bio/crispr-analysis). Cas9 transgenes were segregated out in the T2 generation to stabilize the lines. We analyzed the following homozygous mutants: *chrev1-1, chrev2-2, rev2-5/6, chrev1-1 chrev2-1, chrev1-1 chrev2-7/14, chphb-2, chphb-2 chphv-1, chrev1-1 chcna-1, chrev1-1 chcna-3, chrev1-1 chcna-1 chphb-1, chrev1-1 chcna-1 chphv-1, chrev1-1 chcna-2 chphv-1, chrev1-1 chcna-2 chhb8-1* and *chrev1-1 chcna-1 chphb-1 chphv-1* and corresponding segregating lines. Mutant analysis was performed in the T2 generation except for heteroallelic *chrev2-5/6 and −7/14* mutants in the T1. All mutations and corresponding coding sequences are listed in Fig. S3.

### Light and Dexamethasone (DEX) treatments

#### Light treatment

*C. chenopodiifolia* were grown in specifically designed modules to allow exposure of subterranean fruit to light (light source from the top, fluence rate 85.98 µmol m^-^² s^-1^) while keeping the root system mostly in dark conditions. Transparent 50 mL falcon tubes were cut at the tip to provide an entry of 2 cm diameter. Tubes were then covered with light-proof black cloth and aluminum foil, filled with soil, turned upside down and planted into a pot full of soil (Fig. S1A). Seven-day-old seedlings, previously germinated on plates, were sown in the falcon tubes and watered regularly with a pipette during the first two weeks to prevent drying. Later on, pots were watered using a capillary watering mat. Unless stated otherwise, light treatments were performed by removing the light-proof covering when the first subterranean flower buds reached the soil (around 4 weeks after germination), while the covering was retained in control plants. Fruit at stage 17b were collected after 2 weeks of exposure to light. Only fruit that were growing in contact to the falcon tube wall and clearly exposed to light were collected. Fruits were processed as described.

#### DEX treatment

Four-week-old *C. hirsuta* plants expressing the constructs *ChPER66::GR-Lh4G>>OP4::ChREV1r-GFP* or *ChUBQ10::GR-Lh4G>>OP4::ChMIR165A* were used for DEX induction. Fruit at stage 15 (5 mm length) were marked and all older fruit removed. The prepared stem was then immersed for 10 seconds in solutions of 100, 10, 1 or 0.1 µM water-soluble DEX (Sigma-Aldrich), 0.1% Silwet L-77 or mock (0.1% Silwet L-77) as control. Treatments were repeated daily for 7 consecutive days, then fruit harvested and fixed as described.

#### Histochemistry - Fixation and staining

To visualize SCWs by confocal laser scanning microscopy, cross-sections of the following samples were fixed and stained with an adapted Clearsee protocol^13,72^: fruit valves, fruit, inflorescences and stems. Briefly, samples were embedded in 5% Top vision low melting point agarose (Thermo Scientific) in a 1.5ml Eppendorf tube. Agar blocks were cut in 100-150 µm transverse sections using a Leica Vibratome VT1000 S. Sections were immediately fixed for 1-2 hours at room temperature in 4% paraformaldehyde (PFA, Electron Microscopy Sciences) in PBS solution, washed twice for 1 min with PBS, and cleared in Clearsee solution for at least 24 h with mild shaking. For surface views of the end*b*, fruit valves were dissected and directly fixed in 4% PFA with vacuum using the same process. Valves were cleared until they become transparent. Sections and valves were stained for cellulose with 0.1% Calcofluor White and for lignin with 0.2% Basic Fuchsin in Clearsee solution and incubated overnight. The staining solution was removed and samples rinsed once in fresh Clearsee solution, successively washed for 30 min and 2 hours in fresh Clearsee solution with gentle shaking before mounting in Clearsee for imaging. When only cellulose staining was required, samples were stained in 0.1% Calcofluor white for 30 min to 1 h, then washed as described.

#### Confocal laser scanning microscopy (CLSM)

CLSM was performed on a Leica TCS SP8 with HC PL FLUOTAR (10x/0.30 dry), HC PL APO CS2 (40x/1.10 WATER), HC FLUOTAR L (25x/0.95 WATER) and HCX PL APO lambda blue (63x/1.20 WATER) objectives or on a Leica Stellaris 5 with HC PL FLUOTAR (10x/0.30 dry), HC PL APO CS2 (40x/1.10 WATER) and HC PL APO CS2 (63x/1.30 GLYC). Excitation and detection windows were set as follows: visualization of cellulose/calcofluor white (405 nm, 425–475 nm), lignin/basic fuchsin (561 nm, 600–650 nm), GFP and mNeonGreen (488nm, 490-530nm), mVenus (514nm, 520-550nm). Images were processed using FIJI software^73^.

#### Photography

Photographs of plants, fruit and fruit valves were taken with a Nikon D800 equipped with either an AF-S Micro NIKKOR 105 mm 1: 2.8 G ED or AF-S NIKKOR 24–85 mm 1: 3.5–4.5 G objectives and a Canon D60 with a Canon Macro lens EF-S 35 mm 1:2.8 IS STM objective.

#### Cell wall composition

Fruit valves were collected from mature fruits (two batches of 10 plants) and flash frozen in liquid nitrogen before being freeze-dried and aliquoted into 5 technical replicates per genotype (∼254 mg and 219mg total dry weight for aerial and underground fruit type, respectively). Cellulose content and matrix polysaccharide-derived monosaccharide composition including uronic acids of the samples were determined as described^74,75^. In short, Alcohol Insoluble Residue (AIR) was prepared from freeze-dried samples and split into 2 samples. One half of the samples were treated with a weak acid (4% sulfuric acid) to release matrix polysaccharide-derived monosaccharides, while the other half of the samples were treated initially with a strong acid (72% sulfuric acid) followed by diluting the sulfuric acid concentration to 4% to yield monosaccharides both derived from cellulose and the matrix polymers. Subtraction of the 2 values allows for the quantification of crystalline cellulose. Monosaccharides of all fractions were quantified using an IC Vario high-performance anion-exchange chromatography system 1,068 (Metrohm, Herisau, Switzerland) equipped with a CarboPac PA20 column (Thermo Fisher Scientific, Waltham, MA, USA) and an amperometric detector (Metrohm) using a sodium hydroxide gradient. Acetyl-bromide lignin content of the AIR samples was determined as previously described^76^.

### RNA and miRNA sequencing and analyses

#### Plant material

##### Light induction

*C. chenopodiifolia* plants were grown in falcon tubes as described with subterranean fruit kept in dark conditions for approximatively 6 weeks. At time point 0, light-proof covers were removed from light-treated samples. Subterranean fruit at stage 17ab (initiation of end*b* SCW lignification) were marked. Valves from these fruits were collected after 30 min and 72 h (Fig. 1D). Both valves were carefully dissected from each fruit, taking care that no seeds, replum or septum tissue was included, and immediately flash-frozen in liquid nitrogen. Equivalent dark-treated samples were collected in parallel in the dark using a dim green safe light (light covered with several layers of LEE Filters number 089, fluence rate 0.09 µmol m^-^² s^-1^, most dominant wavelength 523nm). For each sample, ten fruit valves collected from five different plants (one fruit/plant) were pooled together. The experiment was performed in triplicate on three different days at the same hour. Additional samples were collected at t=0 and t=72h for fixation and imaging (Fig. S1B). Samples were used for both long-read and short-read sequencing.

##### DEX treatment

Six-week-old *C. hirsuta* plants expressing the DEX-inducible construct *ChPER66::GR-Lh4G>>OP4::ChREV1r-GFP* were used for RNA-seq. Fruit at stage 15 (5 mm length) were marked the evening before the treatment and all older fruit removed. The next morning, fruit were immersed for 10 seconds in a solution of either 100 µM DEX, 0.1% Silwet L-77 or mock (0.1% Silwet L-77) at time point 0. The treatment was repeated every day at the same hour for four and six consecutive days (Fig. 4B), then valves were dissected as previously described and immediately flash frozen in liquid N_2_. Each sample consisted of a pool of twelve valves coming from three different plants (two fruit/plant). The experiment was performed in triplicate. Samples were used for Illumina short-read sequencing.

##### Arabidopsis and C. hirsuta fruit valves

5.5 to 6.5-week-old *C. hirsuta* plants and 6.5-week-old Arabidopsis plants grown in parallel, in the same conditions, were used to sample fruit valves at stages 16 and 17ab for both total RNA and small RNA sequencing. Plants were carefully staged to reduce variability. Valves were dissected under a binocular lens as previously described and immediately flash frozen in liquid N_2_. Samples consisted of either sixteen valves at stage 17ab or thirty-two valves at stage 16 from eight different plants (i.e. one stage 17ab and two stage 16 fruits harvested from each plant). Arabidopsis samples used for small RNA sequencing consisted of ten stage 17ab and twenty stage 16 valves from five plants. The experiment was carried out separately in quadruplicate for both total RNA and small RNA sequencing. Samples were used for Illumina-short read sequencing.

### Total RNA and small RNA extraction

Frozen tissue was ground using a TissueLyzer with three metal beads per tube, then total RNA extracted with SpectrumTM Plant Total RNA kit (Sigma-Aldrich) following manufacturer’s instruction. An additional DNase I digestion step was included and RNA was eluted in nuclease-free ddH_2_O before further processing. For small RNA sequencing, functional small RNAs from RNA-induced silencing complexes (RISC) were extracted using the TraPR Small RNA Isolation kit (Lexogen) following manufacturer’s instructions. RNA was quantified by spectrophotometry (Nanodrop, Thermo Scientific, USA) and analyzed by capillary electrophoresis using the Agilent 2100 Bioanalyzer (RNA Nanochip, Agilent Technologies, Germany).

### Sequencing

#### Total RNA cDNA library preparation and short-read Illumina sequencing

Total RNA samples from fruit valves of Arabidopsis, *C. hirsuta*, *C. chenopodiifolia* light experiment and *C. hirsuta* DEX experiment, were processed similarly for short-read sequencing. cDNA synthesis, stranded Poly-A selection library preparation (two-sided, 150 bp) and sequencing were carried out by Novogene using the Illumina NovaSeq6000 platform. Raw sequence data are available in ENA bioprojects PRJEB69676 and PRJEB102376.

#### miRNA library preparation and short-read Illumina sequencing

Libraries for small RNA samples from *C. hirsuta* and Arabidopsis fruit valves were prepared by the Max Planck Institute for Plant Breeding Research Genome Centre using the RealSeq®-AC miRNA Library Kit (NextTera Libraries, 500 00048).

Two-sided, 150bp-sequencing was carried out by Novogene as described. Raw sequence data are available in ENA bioproject PRJEB102376.

### Analysis of differentially expressed genes

#### Alignment to C. chenopodiifolia transcriptome

Paired-end short-reads were quality-checked, mapped to the *C. chenopodiifolia* long read reference transcriptome^13^ and transcript abundance quantified using default settings of SALMON v1.8.0^77^. Abundance estimates were imported and transformed to gene level counts using the tximport software^78^. Differential expression analysis was performed with EdgeR (v.3.42.4^79,80^) to test for differences between the four different groups (light-30min, light-72h, dark-30min and dark-72h) using the design formula: Design <-model.matrix (∼0+group). Contrasts between sample groups were done for five comparisons to test for differential expression: light VS dark at 30min and 72h, respectively; 30min VS 72h in light and dark, respectively; and the interaction of light treatment and time. The general linear model Quasilikelihood F-tests (glmQLFTreat) was then used to find significantly differentially expressed genes (DEGs) between the groups. DEGs were selected based on a cut-off with a fold-change (FC) of 2, and a false-discovery rate (FDR) of 0.05.

#### Alignment to C. hirsuta and Arabidopsis genome

Short-reads were quality controlled with FastQC and trimmed using the Cutadapt software^81^ with default values, then aligned to *C. hirsuta*^82^ and Arabidopsis TAIR10 genomes using HISAT2 2.1.0 (default values using the command “hisat2 -p 2 –dta”^83^). Samtools and Qualimap were used for conversion of BAM files and quality control before read count analysis with HTSeq^84–86^. DEG analysis was carried out in R (v.4.2.1) using R studio (v.2022.07.1 +554) and the DESeq2 package v.1.36.0^87^. Design was defined by the formula: design = ∼Stage for comparison of fruit development in *C. hirsuta* and *A. thaliana*. For DEX-induction of *ChREVr,* we used the formula design = ∼ Treatment + Time + Treatment:Time. DEGs were selected based on FC >2 and adjusted p-value < 0.05.

### Cluster and Gene Ontology (GO) analysis

GO annotations previously retrieved for *C. chenopodiifolia* transcripts^13^ and Arabidopsis GO term annotations for orthologous *C. hirsuta* genes^82^ were used to carry out analyses in R with the packages CLUSTERPROFILER v.3.8 for GO analysis^88^, ggplot2 v.3.5.1 for graphs and volcano plots generation^89^, pheatmap v.1.0.12 for heat maps^90^ and VennDiagram v.1.7.3 for Venn diagrams of DEG and reads counts^91^.

### miRNA analysis

Short-reads were quality controlled with FastQC and trimmed using the Cutadapt software with the following settings: -a “TGGAATTCTCGGGTGCCAAGGAACTCCAGTCAC” -A “TGATCGTCGGACTGTAGAACTCTGAACGTGTAGATCTCGGTGGTCGCCGTATCAT” -u 1 –minimum-length 18 – maximum-length 26. Two-sided reads were merged using bbmerge. For Arabidopsis, merged trimmed reads were processed with sRNAbench (sRNAtoolbox^92^). Reads were mapped to the Arabidopsis TAIR10 genome using Bowtie under the default sRNAbench parameters for miRNA profiling (no mismatch allowed). miRNA annotations were taken from miRBase v22.1. Differential expression was performed using sRNAde. For *C. hirsuta*, adapter-trimmed and merged reads were analyzed using Shortstack v. 4.1.2 with the *C. hirsuta* genome^82^ and Arabidopsis mature miRNA annotation from miRBase. Differential expression was performed using DESeq2 v.1.36.0^87^. CPM values for canonical mature miRNA were computed from raw counts.

### Phylogenetic analysis and protein alignment

*HD-ZIPIII* genes were identified in *C. chenopodiifolia* by retrieving *CcREV*, *CcPHB*, *CcPHV*, *CcCNA* and *CcHB-8* sequences from the annotated *C. chenopodiifolia* long read reference transcriptome^13^ and subsequent BLAST search of the *C. chenopodiifolia* genome assembly^14^. In this way, we found sequences for *CcREV2-B* and *C*, *CcHB8-C* and *D* that were present in the *C. chenopodiifolia* genome, but absent from the reference transcriptome. In addition, we also retrieved all sequences belonging to five HD-ZIPIII orthogroups in *C. chenopodiifolia* using OrthoFinder v2.5.5 as previously described^14^. We analyzed gene models by aligning all *C. chenopodiifolia* sequences with *C. hirsuta HD-ZIPIII* CDS sequences using MUSCLE in the Molecular Evolutionary Genetics Analysis (*MEGA12*) software^93^. Protein sequence similarity was calculated using BLOSUM62 matrix with threshold 0. To construct gene trees, *C. chenopodiifolia* CDS were first translated into protein sequences and aligned with HD-ZIPIII sequences from *A. thaliana*, *C. hirsuta* and *Capsella rubella* using MUSCLE. A phylogeny was inferred using the Maximum likelihood method and Jones-Taylor-Thornton (JTT) model of amino acid substitution with rate heterogeneity. The evolutionary rate differences among sites were modeled using a discrete Gamma distribution across four categories. Protein alignments were visualized with Jalview 2.11.4.1.

### Synteny analysis

Intervals containing five genes either side of *REV1* and *REV2* in the diploid *C. hirsuta* genome and the four sub-genomes of *C. chenopodiifolia* were extracted from previous data^14^ generated with the GENESPACE v1.3.1 pipeline^94^ and plotted using R.

### Promoter analysis

Promoter sequences for all DE down-regulated transcripts in common for the interaction fruit stage-fruit type, the comparison aerial vs underground fruit at stage 17ab and the comparison light vs dark at 72h were extracted. Briefly, 3 kb upstream from the start of the transcript or shorter if it overlaps with another upstream gene, was used for further analysis. We used AME (MEME Suite) to identify relatively enriched motifs (JASPAR CORE Plants and ARABIDOPSIS as motif database) in the promoter sequences compared to shuffled promoter sequences (Average odds score and Fisher’s exact test)^95^.

For *REVOLUTA* regulatory sequence analysis, sequences spanning from the first upstream gene (around 10 kb including the 5’UTR) to 5 kb downstream of the stop codon were used for alignment with mVista with the MLAGAN-SHUFFLE alignment program^96^. Conserved non-coding sequences for *C. hirsuta* were extracted from the Conservatory database.

## QUANTIFICATION AND STATISTICAL ANALYSIS

### Quantifications

#### Fruit size and seed number

Stage 17b fruit grown in light or dark as previously described were collected (n=6 fruit from different plants) from three independent experiments. Fruit width and length were measured from images using FIJI software, while pedicel length was measured with a ruler. Seeds were counted manually.

#### Endb SCW polarity

Fruit were sectioned, fixed, stained and imaged by CSLM as previously described. The thickness of the adaxial and abaxial lignified SCWs was measured using FIJI software and the polarity ratio calculated as *polarity ratio = abaxial thickness / adaxial thickness. Light treatment on C.chenopodiifolia fruit:* n = 175, 5 end*b* SCWs in 5-6 valves per treatment in 3 replicates. *DEX-inducible REV1r expression*: n = 100, 10 end*b* SCWs in 1-2 valves from independent plants per treatment, in 5 replicates (one T1 lines and two independent T2 lines) *ChLAC11::ChMYB46-GFP:* n = 55, 9-14 end*b* SCW by valves from 2-3 plants per treatment.

### Valve curvature

Fruit valves were peeled, imaged and the radius (*r*) of the coiled valve was measured using FIJI software. Curvature was calculated as *curvature (K) = 1/radius (r)*.

### Proportion of lignified endb surface

CLSM Z-stacks of the adaxial surface of the end*b* layer (stage 17b fruit) stained for cellulose and lignin were imaged with HC PL FLUOTAR (10x/0.30 dry) and a maximum-projection applied. For *C. hirsuta* fruit valves, the proportion of the end*b* layer that is lignified was measured with a semi-automated script in FIJI. Briefly, the total valve surface was measured on the cell wall channel after application of a Gaussian blurr (radius = 2), automated thresholding using the “Triangle threshold” method and conversion to mask. The threshold was manually curated and adjusted so that the full valve surface was selected. The lignified surface was measured similarly on the raw lignin channel with the “Intermode threshold” and manually curated. The percentage of lignin coverage was calculated as the ratio of lignified surface / total surface. Quantification was made on valve images from genotyped T2 or T3 individual plants (one fruit picture per plant). In total, 157 valves were quantified, with 4 to 23 plants per genotype. For Arabidopsis, most images had low contrast in the lignin channel due to less lignin content in end*b* SCWs compared to *C. hirsuta*. However, lignified end*b* cells also had a stronger cellulose staining than non-lignified cells. We therefore measured the end*b* SCW surface on the raw cellulose channel with the “Default threshold”, and manually curated the threshold. Quantifications of *rev-5* complemented with *AtREV::AtREV-mNG* (n=28 independent T1 lines, 1 fruit/plant), *AtREV::ChREV1-mNG* (n=116, 1 fruit/plant and 3 to 6 plants/line from 18 independent T2 lines) and *AtREV::CcREV1-D-mNG* (n=87, 1 fruit/plant and 3 to 6 plants/line from 15 independent T2 lines) were done in different batches with always wild-type (n=21) and *rev-5* (n=22) samples as control. Control samples with slightly damaged valves that impeded the use of the calcofluor channel for quantification were included in Set1. These samples were quantified as for *C. hirsuta* since they had strong Basic fuchsin staining.

### Complementation of rev-5 phenotypes in Arabidopsis

The same complementation lines described above were assessed for floral and axillary meristem phenotypes. Floral meristem defects were scored as “non-developing” fruit (i.e. filamentous structures typical of *rev-5*, or sterile flowers that lack carpels and left only a pedicel after floral organ abscission) in the first 20 fruits, and the ratio of normal fruit/total fruit was calculated. On the same plants, axillary meristem defects were scored as the ratio of secondary cauline leaves developing a 3^rd^ lateral shoot meristem (LSM) as follows: all cauline leaves of the secondary branches were scored for presence or absence of a tertiary LSM and the ratio calculated as number of leaves with 3^rd^ LSM/total number of secondary cauline leaves. Sample numbers: wild-type (n = 22), *rev-*5 (n = 26), *rev-5* complemented with *AtREV::AtREV-mNG* (n=28), *AtREV::ChREV1-mNG* (n=135) and *AtREV::CcREV1-D-mNG* (n=122).

Complementation of *rev-5* was also assessed by crossing with *REV::REV-mVenus*. Among 53 T2 progeny, 4 plants exhibited the *rev-5* floral meristem phenotype. Based on an expected ratio of 1/16 for complemented plants, the Pearson’s chi-square value was 0.153, which is below the critical value (3.84; *df* = 1, α = 0.05), consistent with full complementation of the *rev-5* phenotype by *REV::REV-mVenus*.

### Statistical analysis

All statistical analyses were carried out using R (v.4.2.1)^97^ and RStudio (2022.07.1+554)^98^ with the following packages: stats v.4.2.1., rstatix v.0.7.2, dplyr v.1.1.4, ggplot2 v.3.5.2, tidyr v.1.3.1^89,99–101^. Statistical details of experiments, including sample number (*n*), number of replicates, and statistical tests, are reported in figure legends and methods. Boxplots display the median (thick line), interquartile range (IQR, box) and 1.5 times the IQR (whiskers). Binary comparisons were performed using Student t-test. When the data did not follow the linear model assumption (assessed using Shapiro and Levene tests), Wilcoxon Mann-Whitney test was used for binary comparisons. Statistically significant differences between groups were indicated by asterisks (P-values: ≥ 0.05 [n.s.], < 0.05 *, < 0.01 **, < 0.001 ***). For multiple comparisons, ANOVA followed by the Tukey’s Honestly Significant difference (HSD) test were applied when linear model assumptions were met. When the data did not meet linear model assumptions, a Kruskal-Wallis test followed by Dunn’s multiple comparison test was performed. Statistically significant differences between groups were indicated by non-shared letters (P < 0.05 unless stated otherwise). Pearson correlations were calculated using the package stats v.4.2.1.

## KEY RESOURCES TABLE

| REAGENT or RESOURCE | SOURCE | IDENTIFIER |
| --- | --- | --- |
| Bacterial and virus strains |  |  |
| *Agrobacterium tumefaciens* GV3101 | N/A | N/A |
| *Escherichia coli* DH10B | N/A | N/A |
| Chemicals, peptides, and recombinant proteins |  |  |
| Dexamethasone-Water Soluble | Sigma-Aldrich | Cat#D2915 |
| Top vision low melting point agarose | Thermo Scientific | Cat#R0801 |
| Paraformaldehyde 20% Solution, EM Grade | Electron Microscopy Sciences | Cat#15713S |
| Calcofluor White / Fluorescent Brightener 28 | Sigma-Aldrich | Cat#910090 |
| Basic Fuchsin | Sigma-Aldrich | Cat#857343 |
| DNase I | Sigma-Aldrich | Cat#DNASE70 |
| Critical commercial assays |  |  |
| In-Fusion Advantage PCR cloning | Clontech | Cat#638956 |
| MultiSite Gateway™ Pro Plus | Invitrogen, Thermo Fisher Scientific | Cat#12537100 |
| Spectrum™ Plant Total RNA kit | Sigma-Aldrich | Cat#STRN50 |
| TraPR Small RNA Isolation kit | Lexogen | Cat#128.24 |
| RealSeq®-AC miRNA Library Kit | NextTera Libraries | Cat#500 00048 |
| Deposited data |  |  |
| *C. chenopodiifolia* long read reference transcriptome | Emonet et al.<sup>13</sup> | ENA: PRJEB69676<br>https://www.ebi.ac.uk/ena/browser/view/PRJEB69676 |
| *C. chenopodiifolia* genome assembly | Emonet et al.<sup>14</sup> |  |
| *C. chenopodiifolia* Orthofinder data | Emonet et al.<sup>14</sup> | https://doi.org/10.5524/102639 |
| TAIR10 Arabidopsis annotation | TAIR | ftp://ftp.arabidopsis.org/home/tair/Genes/TAIR10_genome_release/ |
| *C. hirsuta* genome assembly/annotation | Gan et al.<sup>82</sup> | PRJNA293154<br>https://www.ncbi.nlm.nih.gov/bioproject/?term=PRJNA293154 |
| *C. chenopodiifolia* RNA sequencing data for fruit | Emonet et al.<sup>13</sup> | ENA: PRJEB69676<br>https://www.ebi.ac.uk/ena/browser/view/PRJEB69676 |
| Raw RNA sequencing data for *C. chenopodiifolia* | This paper, Emonet et al.<sup>13</sup> | ENA: PRJEB69676<br>https://www.ebi.ac.uk/ena/browser/view/PRJEB69676 |
| Raw RNA sequencing data for Arabidopsis and *C. hirsuta* | This paper | ENA: PRJEB102376<br>https://www.ebi.ac.uk/ena/browser/view/PRJEB102376 |
| JASPAR CORE Plants and ARABIDOPSIS | Rauluseviciute et al.<sup>102</sup> | https://jaspar.elixir.no/search?q=&collection=CORE&tax_group=plants |
| Conservatory database | Hendelam et al.<sup>50</sup> | GSE164297 |
| Experimental models: Organisms/strains |  |  |
| Arabidopsis: wild-type / Col-0 | N/A | N/A |
| *Cardamine hirsuta*: wild type / Ox | Hay et al.<sup>62</sup> | Herbarium specimen voucher Hay 1 (OXF) |
| *Cardamine chenopodiifolia* Pers. (Ipen: XX-0-MJG-19—35600) | Emonet et al.<sup>13</sup> | N/A |
| *Arabidopsis*: *rev-5*/ Col-0 | Talbert et al.<sup>43</sup> | N/A |
| *Arabidopsis*: *rev-9*/ Col-0 | Emery et al.<sup>30</sup> | N/A |
| *Arabidopsis*: *rev-10D*/ Col-0 | Emery et al.<sup>30</sup> | N/A |
| *Arabidopsis*: *phb-13 phv-11* / Col-0 | Prigge et al.<sup>42</sup> | N/A |
| *Arabidopsis*: *phb phv* / Col-0 | Kim et al.<sup>63</sup> | N/A |
| *Arabidopsis*: *phb phv rev-5 +/-* / Col-0 | Dello Ioio et al.<sup>41</sup> | N/A |
| *Arabidopsis*: *phb-13 phv-11 cna-2* / Col-0 | Prigge et al.<sup>42</sup> | N/A |
| *Arabidopsis*: *phb-13 phv-11 cna-2 athb8-11*/ Col-0 | Prigge et al.<sup>42</sup> | N/A |
| *Arabidopsis*: *MIR166A::erGFP*/ Col-0 | Burian et al.<sup>40</sup> | N/A |
| *Arabidopsis*: *REV::REV-VENUS* / Col-0 | Heisler et al.<sup>34</sup> | N/A |
| *Arabidopsis*: *Cre-U5-Cre_CAB3::lox-CyPET-lox-3xVenus-RCI2A (CAB3::3xVenus-RCI2A )*/ Col-0 | C. Whitewoods | N/A |
| *Arabidopsis*: *CAB3::3xVenus-RCI2A rev-5* / Col-0 | This paper | N/A |
| *Arabidopsis*: *REV::REV-VENUS rev-5* / Col-0 | This paper | N/A |
| *Arabidopsis*: *MIR166A::erGFP rev-5*/ Col-0 | This paper | N/A |
| *Arabidopsis*: *AtREV::AtREV-mNG rev-5*/ Col-0 | This paper | N/A |
| *Arabidopsis*: *AtREV::ChREV1::mNG rev-5*/ Col-0 | This paper | N/A |
| *Arabidopsis*: *AtREV::CcREV1-D-mNG rev-5*/ Col-0 | This paper | N/A |
| *C. hirsuta*: *chrev1-1* / Ox | This paper | N/A |
| *C. hirsuta*: *chrev2-2*/ Ox | This paper | N/A |
| *C. hirsuta*: *chrev2-5/6*/ Ox | This paper | N/A |
| *C. hirsuta*: *chrev1-1 chrev2-1*/ Ox | This paper | N/A |
| *C. hirsuta*: *chrev1-1 chrev2-7/14*/ Ox | This paper | N/A |
| *C. hirsuta*: *chphb-2*/ Ox | This paper | N/A |
| *C. hirsuta*: *chphb-2 chphv-1*/ Ox | This paper | N/A |
| *C. hirsuta*: *chrev1-1 chcna-1*/ Ox | This paper | N/A |
| *C. hirsuta*: *chrev1-1 chcna-3*/ Ox | This paper | N/A |
| *C. hirsuta*: *chrev1-1 chcna-1 chphb-1*/ Ox | This paper | N/A |
| *C. hirsuta*: *chrev1-1 chcna-1 chphv-1*/ Ox | This paper | N/A |
| *C. hirsuta*: *chrev1-1 chcna-2 chphv-1*/ Ox | This paper | N/A |
| *C. hirsuta*: *chrev1-1 chcna-2 chhb8-1* / Ox | This paper | N/A |
| *C. hirsuta*: *chrev1-1 chcna-1 chphb-1 chphv-1* / Ox | This paper | N/A |
| *C. hirsuta*: *ChREV1::ChREV1-mNG*/ Ox | This paper | N/A |
| *C. hirsuta* : *ChREV1::ChREV1-mNG chrev1-1*/ Ox | This paper | N/A |
| *C. hirsuta*: *ChPER66::GR-LhG4>>pOP4::ChREV1r-GFP*/ Ox | This paper | N/A |
| *C. hirsuta*: *ChUBQ10::GR-LhG4>>OP4::ChMIR165A*/ Ox | This paper | N/A |
| *C. hirsuta*: *ChLAC11::ChMYB46-GFP*/ Ox | This paper | N/A |
| *C. hirsuta*: *ChMYB46::NLS-3xGFP*/ Ox | This paper | N/A |
| Oligonucleotides |  |  |
| Primers for genotyping and sgRNA listed in Table S3 | Microsynth, Sigma | N/A |
| Recombinant DNA |  |  |
| Multisite GreenGate vectors | Lampropoulos et al.<sup>64</sup> | N/A |
| pGGD mNeonGreen | Eng et al.<sup>11</sup> | N/A |
| *pGGA-pChPER66* | Perez-Anton et al.<sup>12</sup> | N/A |
| *pGGA-pChLAC11* | Perez-Anton et al.<sup>12</sup> | N/A |
| pGGZwf01 | Perez-Anton et al.<sup>12</sup> | N/A |
| *ChLAC11::ChMYB46c-GFP* | This paper | N/A |
| *ChMYB46::NLS-3xGFP* | This paper | N/A |
| *ChREV1::ChREV1-mNG* | This paper | N/A |
| *AtREV::AtREV-mNG* | This paper | N/A |
| *AtREV::ChREV1-mNG* | This paper | N/A |
| *AtREV::CcREV1-D-mNG* | This paper | N/A |
| *ChPER66::GR-Lh4G>>OP4::ChREV1r-GFP* | This paper | N/A |
| *ChUBQ10::GR-Lh4G>>OP4::ChMIR165A* | This paper | N/A |
| *pOP4-pGGA* | Schurholz et al.<sup>66</sup> | N/A |
| *GR-LhG4-pGGC* | Schurholz et al.<sup>66</sup> | N/A |
| pDeCas9 | Fauser et al.<sup>68</sup> | N/A |
| pRU41-pRU48 | Ursache et al.<sup>69</sup> | N/A |
| pSF464 | Ursache et al.<sup>69</sup> | N/A |
| pRU325 | Ursache et al.<sup>69</sup> | N/A |
| pRU295 *pEC1.2::zCas9i* FastGreen | Ursache et al.<sup>69</sup> | N/A |
| pRU294 *pEC1.2::zCas9i* FastRed | Ursache et al.<sup>69</sup> | N/A |
| Software and algorithms |  |  |
| CCTop | Stemmer et al.<sup>70</sup> | https://cctop.cos.uni-heidelberg.de |
| CRISPRater | Labuhn et al.<sup>71</sup> | https://cctop.cos.uni-heidelberg.de |
| Geneious Prime v.10.2.6 | Biomatters Ltd | http://www.geneious.com |
| ICE CRISPR Analysis Tool | EditCo | https://www.editco.bio/crispr-analysis |
| FIJI/ImageJ 1.54p | Schindelin et al.<sup>73</sup> | N/A |
| SALMON v1.8.0 | Patro et al.<sup>77</sup> | N/A |
| tximport | Soneson et al.<sup>78</sup> | N/A |
| EdgeR v.3.42.4 | Robinson et al.<sup>79,80</sup> | N/A |
| FastQC v0.12.1 | Babraham Bioinformatics | https://www.bioinformatics.babraham.ac.uk/projects/fastqc/ |
| Cutadapt v.5.1 | Martin et al.<sup>81</sup> | N/A |
| HISAT2 v2.1.0 | Kim et al.<sup>83</sup> | N/A |
| Samtools v.1.21 | Danecek et al.<sup>86</sup> | N/A |
| Qualimap v.2.2.1 | Garcia-Alcalde et al.<sup>84</sup> | N/A |
| HTSeq 0.6.1p1 | Anders et al.<sup>85</sup> | N/A |
| R v.4.2.1 | R Core Team<sup>97</sup> | https://www.R-project.org/ |
| R studio v.2022.07.1 +554 | Posit Team<sup>98</sup> | http://www.rstudio.com/ |
| DESeq2 v.1.36.0 | Love et al.<sup>87</sup> | N/A |
| CLUSTERPROFILER v.3.8 | Wu et al.<sup>88</sup> | N/A |
| ggplot2 v.3.5.1 | Wickham et al.<sup>89</sup> | N/A |
| pheatmap v.1.0.12 | Kolde et al.<sup>90</sup> | N/A |
| VennDiagram v.1.7.3 | Chen et al.<sup>91</sup> | N/A |
| blast 2.14.1 | Camacho et al.<sup>103</sup> | N/A |
| MEGA12 v.12.0.11 | Kumar et al.<sup>93</sup> | N/A |
| MUSCLE v3.8.31 | Edgar et al.<sup>104</sup> | N/A |
| Jalview 2.11.4.1. | Waterhouse et al.<sup>105</sup> | N/A |
| GENESPACE v1.3.1 | Lovell et al.<sup>94</sup> | N/A |
| AME (MEME Suite) | McLeay et al.<sup>95</sup> | N/A |
| mVista | Brudno et al.<sup>96</sup> | N/A |
| stats v.4.2.1., rstatix v.0.7.2 | Kassambara et al.<sup>99</sup> | N/A |
| dplyr v.1.1.4 | Wickham et al.<sup>100</sup> | N/A |
| tidyr v.1.3.1 | Wickham et al.<sup>101</sup> | N/A |

## References

1. Stuppy, W., and Kesseler, R. (2011). Fruit - Edible, Inedible, Incredible, 2nd Edition (Papadakis).

2. Kokko, H., and Lopez-Sepulcre, A. (2006). From individual dispersal to species ranges: perspectives for a changing world. Science 313, 789–791. 10.1126/science.1128566.

3. Hamilton, W.D., and May, R.M. (1977). Dispersal in Stable Habitats. Nature 269, 578–581. Doi 10.1038/269578a0.

4. Emonet, A., and Hay, A. (2024). Explosive seed dispersal. Curr Biol 34, R970–R972. 10.1016/j.cub.2024.07.050.

5. Hofhuis, H., Moulton, D., Lessinnes, T., Routier-Kierzkowska, A.L., Bomphrey, R.J., Mosca, G., Reinhardt, H., Sarchet, P., Gan, X., Tsiantis, M., et al. (2016). Morphomechanical Innovation Drives Explosive Seed Dispersal. Cell 166, 222–233. 10.1016/j.cell.2016.05.002.

6. Mosca, G., Eng, R.C., Adibi, M., Yoshida, S., Lane, B., Bergheim, L., Weber, G., Smith, R.S., and Hay, A. (2024). Growth and tension in explosive fruit. Curr Biol 34, 1010–1022 e1014. 10.1016/j.cub.2024.01.059.

7. Spence, J., Vercher, Y., Gates, P., and Harris, N. (1996). ’Pod shatter’ in *Arabidopsis thaliana*, Brassica napus and B-juncea. J Microsc-Oxford 181, 195–203. DOI 10.1046/j.1365-2818.1996.111391.x.

8. Mitsuda, N., and Ohme-Takagi, M. (2008). NAC transcription factors NST1 and NST3 regulate pod shattering in a partially redundant manner by promoting secondary wall formation after the establishment of tissue identity. Plant J 56, 768–778. 10.1111/j.1365-313X.2008.03633.x.

9. Kumar, M., Campbell, L., and Turner, S. (2016). Secondary cell walls: biosynthesis and manipulation. J Exp Bot 67, 515–531. 10.1093/jxb/erv533.

10. Taylor-Teeples, M., Lin, L., de Lucas, M., Turco, G., Toal, T.W., Gaudinier, A., Young, N.F., Trabucco, G.M., Veling, M.T., Lamothe, R., et al. (2015). An Arabidopsis gene regulatory network for secondary cell wall synthesis. Nature 517, 571–575. 10.1038/nature14099.

11. Eng, R.C., Emonet, A., Neumann, U., Pauly, M., and Hay, A. (2026). CESA7 and microtubules pattern complex secondary cell walls in explosive fruit of *Cardamine hirsuta*. Plant Cell 38. 10.1093/plcell/koag062.

12. Perez-Anton, M., Schneider, I., Kroll, P., Hofhuis, H., Metzger, S., Pauly, M., and Hay, A. (2022). Explosive seed dispersal depends on SPL7 to ensure sufficient copper for localized lignin deposition via laccases. Proc Natl Acad Sci U S A 119, e2202287119. 10.1073/pnas.2202287119.

13. Emonet, A., Perez-Anton, M., Neumann, U., Dunemann, S., Huettel, B., Koller, R., and Hay, A. (2024). Amphicarpic development in *Cardamine chenopodiifolia*. New Phytol 244, 1041–1056. 10.1111/nph.19965.

14. Emonet, A., Awad, M., Tikhomirov, N., Vasilarou, M., Perez-Anton, M., Gan, X., Novikova, P.Y., and Hay, A. (2024). Polyploid genome assembly of *Cardamine chenopodiifolia*. GigaByte 2024, gigabyte145. 10.46471/gigabyte.145.

15. Gianella, M., Bradford, K.J., and Guzzon, F. (2021). Ecological, (epi)genetic and physiological aspects of bet-hedging in angiosperms. Plant Reprod 34, 21–36. 10.1007/s00497-020-00402-z.

16. Dearden, P.K. (2025). Polyphenisms: a developmental perspective. Development 152. 10.1242/dev.204693.

17. West-Eberhard, M.J. (2003). Developmental plasticity and evolution (Oxford University Press). 10.1093/oso/9780195122343.001.0001.

18. Moczek, A.P., Sultan, S., Foster, S., Ledon-Rettig, C., Dworkin, I., Nijhout, H.F., Abouheif, E., and Pfennig, D.W. (2011). The role of developmental plasticity in evolutionary innovation. Proceedings. Biological sciences / The Royal Society 278, 2705–2713. 10.1098/rspb.2011.0971.

19. Stebbins, G.L., Jr. (1947). Types of polyploids; their classification and significance. Adv Genet 1, 403–429. 10.1016/s0065-2660(08)60490-3.

20. Ohno, S. (1970). Evolution by Gene Duplication (Springer). 10.1007/978-3-642-86659-3.

21. Holland, P.W., Marletaz, F., Maeso, I., Dunwell, T.L., and Paps, J. (2017). New genes from old: asymmetric divergence of gene duplicates and the evolution of development. Philos Trans R Soc Lond B Biol Sci 372. 10.1098/rstb.2015.0480.

22. Vlad, D., Kierzkowski, D., Rast, M.I., Vuolo, F., Dello Ioio, R., Galinha, C., Gan, X., Hajheidari, M., Hay, A., Smith, R.S., et al. (2014). Leaf shape evolution through duplication, regulatory diversification, and loss of a homeobox gene. Science 343, 780–783. 10.1126/science.1248384.

23. Vuolo, F., Mentink, R.A., Hajheidari, M., Bailey, C.D., Filatov, D.A., and Tsiantis, M. (2016). Coupled enhancer and coding sequence evolution of a homeobox gene shaped leaf diversity. Genes Dev 30, 2370–2375. 10.1101/gad.290684.116.

24. Carabelli, M., Morelli, G., Whitelam, G., and Ruberti, I. (1996). Twilight-zone and canopy shade induction of the *Athb-2* homeobox gene in green plants. Proc Natl Acad Sci U S A 93, 3530– 3535. 10.1073/pnas.93.8.3530.

25. Ruberti, I., Sessa, G., Lucchetti, S., and Morelli, G. (1991). A novel class of plant proteins containing a homeodomain with a closely linked leucine zipper motif. EMBO J 10, 1787–1791. 10.1002/j.1460-2075.1991.tb07703.x.

26. Schena, M., and Davis, R.W. (1992). HD-Zip proteins: members of an Arabidopsis homeodomain protein superfamily. Proc Natl Acad Sci U S A 89, 3894–3898. 10.1073/pnas.89.9.3894.

27. Tepperman, J.M., Zhu, T., Chang, H.S., Wang, X., and Quail, P.H. (2001). Multiple transcription-factor genes are early targets of phytochrome A signaling. Proc Natl Acad Sci U S A 98, 9437– 9442. 10.1073/pnas.161300998.

28. Roig-Villanova, I., Bou, J., Sorin, C., Devlin, P.F., and Martinez-Garcia, J.F. (2006). Identification of primary target genes of phytochrome signaling. Early transcriptional control during shade avoidance responses in Arabidopsis. Plant Physiol 141, 85–96. 10.1104/pp.105.076331.

29. Pfeiffer, A., Shi, H., Tepperman, J.M., Zhang, Y., and Quail, P.H. (2014). Combinatorial complexity in a transcriptionally centered signaling hub in Arabidopsis. Mol Plant 7, 1598– 1618. 10.1093/mp/ssu087.

30. Emery, J.F., Floyd, S.K., Alvarez, J., Eshed, Y., Hawker, N.P., Izhaki, A., Baum, S.F., and Bowman, J.L. (2003). Radial patterning of Arabidopsis shoots by class III HD-ZIP and KANADI genes. Curr Biol 13, 1768–1774. 10.1016/j.cub.2003.09.035.

31. Merelo, P., Ram, H., Pia Caggiano, M., Ohno, C., Ott, F., Straub, D., Graeff, M., Cho, S.K., Yang, S.W., Wenkel, S., and Heisler, M.G. (2016). Regulation of MIR165/166 by class II and class III homeodomain leucine zipper proteins establishes leaf polarity. Proc Natl Acad Sci U S A 113, 11973–11978. 10.1073/pnas.1516110113.

32. Brandt, R., Salla-Martret, M., Bou-Torrent, J., Musielak, T., Stahl, M., Lanz, C., Ott, F., Schmid, M., Greb, T., Schwarz, M., et al. (2012). Genome-wide binding-site analysis of REVOLUTA reveals a link between leaf patterning and light-mediated growth responses. Plant J 72, 31–42. 10.1111/j.1365-313X.2012.05049.x.

33. Reinhart, B.J., Liu, T., Newell, N.R., Magnani, E., Huang, T., Kerstetter, R., Michaels, S., and Barton, M.K. (2013). Establishing a framework for the Ad/abaxial regulatory network of Arabidopsis: ascertaining targets of class III homeodomain leucine zipper and KANADI regulation. Plant Cell 25, 3228–3249. 10.1105/tpc.113.111518.

34. Heisler, M.G., Ohno, C., Das, P., Sieber, P., Reddy, G.V., Long, J.A., and Meyerowitz, E.M. (2005). Patterns of auxin transport and gene expression during primordium development revealed by live imaging of the Arabidopsis inflorescence meristem. Curr Biol 15, 1899–1911. 10.1016/j.cub.2005.09.052.

35. Zhong, R., and Ye, Z.H. (1999). *IFL1*, a gene regulating interfascicular fiber differentiation in Arabidopsis, encodes a homeodomain-leucine zipper protein. Plant Cell 11, 2139–2152. 10.1105/tpc.11.11.2139.

36. McConnell, J.R., Emery, J., Eshed, Y., Bao, N., Bowman, J., and Barton, M.K. (2001). Role of PHABULOSA and PHAVOLUTA in determining radial patterning in shoots. Nature 411, 709–713. 10.1038/35079635.

37. Otsuga, D., DeGuzman, B., Prigge, M.J., Drews, G.N., and Clark, S.E. (2001). REVOLUTA regulates meristem initiation at lateral positions. Plant J 25, 223–236. 10.1111/j.1365-313X.2001.00959.x.

38. Rhoades, M.W., Reinhart, B.J., Lim, L.P., Burge, C.B., Bartel, B., and Bartel, D.P. (2002). Prediction of Plant MicroRNA Targets. Cell 110, 513–520. 10.1016/S0092-8674(02)00863-2.

39. Reinhart, B.J., Weinstein, E.G., Rhoades, M.W., Bartel, B., and Bartel, D.P. (2002). MicroRNAs in plants. Genes Dev 16, 1616–1626. 10.1101/gad.1004402.

40. Burian, A., Paszkiewicz, G., Nguyen, K.T., Meda, S., Raczynska-Szajgin, M., and Timmermans, M.C.P. (2022). Specification of leaf dorsiventrality via a prepatterned binary readout of a uniform auxin input. Nat Plants 8, 269–280. 10.1038/s41477-022-01111-3.

41. Dello Ioio, R., Galinha, C., Fletcher, A.G., Grigg, S.P., Molnar, A., Willemsen, V., Scheres, B., Sabatini, S., Baulcombe, D., Maini, P.K., and Tsiantis, M. (2012). A PHABULOSA/cytokinin feedback loop controls root growth in Arabidopsis. Curr Biol 22, 1699–1704. 10.1016/j.cub.2012.07.005.

42. Prigge, M.J., Otsuga, D., Alonso, J.M., Ecker, J.R., Drews, G.N., and Clark, S.E. (2005). Class III Homeodomain-Leucine Zipper Gene Family Members Have Overlapping, Antagonistic, and Distinct Roles in Arabidopsis Development. Plant Cell 17, 61–76. 10.1105/tpc.104.026161.

43. Talbert, P.B., Adler, H.T., Parks, D.W., and Comai, L. (1995). The REVOLUTA gene is necessary for apical meristem development and for limiting cell divisions in the leaves and stems of *Arabidopsis thaliana*. Development 121, 2723–2735. 10.1242/dev.121.9.2723.

44. Magnani, E., and Barton, M.K. (2011). A per-ARNT-sim-like sensor domain uniquely regulates the activity of the homeodomain leucine zipper transcription factor REVOLUTA in Arabidopsis. Plant Cell 23, 567–582. 10.1105/tpc.110.080754.

45. Husbands, A.Y., Feller, A., Aggarwal, V., Dresden, C.E., Holub, A.S., Ha, T., and Timmermans, M.C.P. (2023). The START domain potentiates HD-ZIPIII transcriptional activity. Plant Cell 35, 2332–2348. 10.1093/plcell/koad058.

46. Kim, W.C., Ko, J.H., and Han, K.H. (2012). Identification of a cis-acting regulatory motif recognized by MYB46, a master transcriptional regulator of secondary wall biosynthesis. Plant Mol Biol 78, 489–501. 10.1007/s11103-012-9880-7.

47. Zhong, R., and Ye, Z.H. (2012). MYB46 and MYB83 bind to the SMRE sites and directly activate a suite of transcription factors and secondary wall biosynthetic genes. Plant Cell Physiol 53, 368–380. 10.1093/pcp/pcr185.

48. Kubo, M., Udagawa, M., Nishikubo, N., Horiguchi, G., Yamaguchi, M., Ito, J., Mimura, T., Fukuda, H., and Demura, T. (2005). Transcription switches for protoxylem and metaxylem vessel formation. Genes Dev 19, 1855–1860. 10.1101/gad.1331305.

49. Mallory, A.C., Reinhart, B.J., Jones-Rhoades, M.W., Tang, G., Zamore, P.D., Barton, M.K., and Bartel, D.P. (2004). MicroRNA control of PHABULOSA in leaf development: importance of pairing to the microRNA 5’ region. The EMBO Journal 23, 3356–3364. 10.1038/sj.emboj.7600340.

50. Hendelman, A., Zebell, S., Rodriguez-Leal, D., Dukler, N., Robitaille, G., Wu, X., Kostyun, J., Tal, L., Wang, P., Bartlett, M.E., et al. (2021). Conserved pleiotropy of an ancient plant homeobox gene uncovered by cis-regulatory dissection. Cell 184, 1724–1739 e1716. 10.1016/j.cell.2021.02.001.

51. Carlsbecker, A., Lee, J.Y., Roberts, C.J., Dettmer, J., Lehesranta, S., Zhou, J., Lindgren, O., Moreno-Risueno, M.A., Vaten, A., Thitamadee, S., et al. (2010). Cell signalling by microRNA165/6 directs gene dose-dependent root cell fate. Nature 465, 316–321. 10.1038/nature08977.

52. Wallner, E.S., and Dolan, L. (2024). Reproducibly oriented cell divisions pattern the prothallus to set up dorsoventrality and de novo meristem formation in Marchantia polymorpha. Curr Biol 34, 4357–4367 e4354. 10.1016/j.cub.2024.07.099.

53. Goethe, J.W.v., and Miller, G.L. (2009). The metamorphosis of plants (MIT Press).

54. Smetana, O., Makila, R., Lyu, M., Amiryousefi, A., Sanchez Rodriguez, F., Wu, M.F., Sole-Gil, A., Leal Gavarron, M., Siligato, R., Miyashima, S., et al. (2019). High levels of auxin signalling define the stem-cell organizer of the vascular cambium. Nature 565, 485–489. 10.1038/s41586-018-0837-0.

55. Miyashima, S., Koi, S., Hashimoto, T., and Nakajima, K. (2011). Non-cell-autonomous microRNA165 acts in a dose-dependent manner to regulate multiple differentiation status in the Arabidopsis root. Development 138, 2303–2313. 10.1242/dev.060491.

56. Leivar, P., and Quail, P.H. (2011). PIFs: pivotal components in a cellular signaling hub. Trends Plant Sci 16, 19–28. 10.1016/j.tplants.2010.08.003.

57. Ni, M., Tepperman, J.M., and Quail, P.H. (1999). Binding of phytochrome B to its nuclear signalling partner PIF3 is reversibly induced by light. Nature 400, 781–784. 10.1038/23500.

58. Al-Sady, B., Ni, W., Kircher, S., Schafer, E., and Quail, P.H. (2006). Photoactivated phytochrome induces rapid PIF3 phosphorylation prior to proteasome-mediated degradation. Mol Cell 23, 439–446. 10.1016/j.molcel.2006.06.011.

59. Juarez, M.T., Kui, J.S., Thomas, J., Heller, B.A., and Timmermans, M.C. (2004). microRNA-mediated repression of rolled leaf1 specifies maize leaf polarity. Nature 428, 84–88.

60. Zhang, T., Li, Y., Ma, L., Sang, X., Ling, Y., Wang, Y., Yu, P., Zhuang, H., Huang, J., Wang, N., et al. (2017). LATERAL FLORET 1 induced the three-florets spikelet in rice. Proc Natl Acad Sci U S A 114, 9984–9989. 10.1073/pnas.1700504114.

61. Dixon, L.E., Pasquariello, M., Badgami, R., Levin, K.A., Poschet, G., Ng, P.Q., Orford, S., Chayut, N., Adamski, N.M., Brinton, J., et al. (2022). MicroRNA-resistant alleles of *HOMEOBOX DOMAIN-2* modify inflorescence branching and increase grain protein content of wheat. Sci Adv 8, eabn5907. 10.1126/sciadv.abn5907.

62. Hay, A., and Tsiantis, M. (2006). The genetic basis for differences in leaf form between *Arabidopsis thaliana* and its wild relative *Cardamine hirsuta*. Nat Genet 38, 942–947. 10.1038/ng1835.

63. Kim, Y.S., Kim, S.G., Lee, M., Lee, I., Park, H.Y., Seo, P.J., Jung, J.H., Kwon, E.J., Suh, S.W., Paek, K.H., and Park, C.M. (2008). HD-ZIP III activity is modulated by competitive inhibitors via a feedback loop in Arabidopsis shoot apical meristem development. Plant Cell 20, 920–933. 10.1105/tpc.107.057448.

64. Lampropoulos, A., Sutikovic, Z., Wenzl, C., Maegele, I., Lohmann, J.U., and Forner, J. (2013). GreenGate---a novel, versatile, and efficient cloning system for plant transgenesis. PLoS One 8, e83043. 10.1371/journal.pone.0083043.

65. Clough, S.J., and Bent, A.F. (1998). Floral dip: a simplified method for Agrobacterium-mediated transformation of *Arabidopsis thaliana*. Plant J 16, 735–743. 10.1046/j.1365-313x.1998.00343.x.

66. Schurholz, A.K., Lopez-Salmeron, V., Li, Z., Forner, J., Wenzl, C., Gaillochet, C., Augustin, S., Barro, A.V., Fuchs, M., Gebert, M., et al. (2018). A Comprehensive Toolkit for Inducible, Cell Type-Specific Gene Expression in Arabidopsis. Plant Physiol 178, 40–53. 10.1104/pp.18.00463.

67. Alvim Kamei, C.L., Pieper, B., Laurent, S., Tsiantis, M., and Huijser, P. (2020). CRISPR/Cas9-Mediated Mutagenesis of *RCO* in *Cardamine hirsuta*. Plants 9, 268. 10.3390/plants9020268.

68. Fauser, F., Schiml, S., and Puchta, H. (2014). Both CRISPR/Cas-based nucleases and nickases can be used efficiently for genome engineering in *Arabidopsis thaliana*. Plant J 79, 348–359. 10.1111/tpj.12554.

69. Ursache, R., Fujita, S., Denervaud Tendon, V., and Geldner, N. (2021). Combined fluorescent seed selection and multiplex CRISPR/Cas9 assembly for fast generation of multiple Arabidopsis mutants. Plant Methods 17, 111. 10.1186/s13007-021-00811-9.

70. Stemmer, M., Thumberger, T., Del Sol Keyer, M., Wittbrodt, J., and Mateo, J.L. (2015). CCTop: An Intuitive, Flexible and Reliable CRISPR/Cas9 Target Prediction Tool. PLoS One 10, e0124633. 10.1371/journal.pone.0124633.

71. Labuhn, M., Adams, F.F., Ng, M., Knoess, S., Schambach, A., Charpentier, E.M., Schwarzer, A., Mateo, J.L., Klusmann, J.H., and Heckl, D. (2018). Refined sgRNA efficacy prediction improves large- and small-scale CRISPR-Cas9 applications. Nucleic Acids Res 46, 1375–1385. 10.1093/nar/gkx1268.

72. Ursache, R., Andersen, T.G., Marhavy, P., and Geldner, N. (2018). A protocol for combining fluorescent proteins with histological stains for diverse cell wall components. Plant J 93, 399–412. 10.1111/tpj.13784.

73. Schindelin, J., Arganda-Carreras, I., Frise, E., Kaynig, V., Longair, M., Pietzsch, T., Preibisch, S., Rueden, C., Saalfeld, S., Schmid, B., et al. (2012). Fiji: an open-source platform for biological-image analysis. Nat Methods 9, 676–682. 10.1038/nmeth.2019.

74. Bauer, S., and Ibanez, A.B. (2014). Rapid determination of cellulose. Biotechnol Bioeng 111, 2355–2357. 10.1002/bit.25276.

75. Yeats, T., Vellosillo, T., Sorek, N., Ibáñez, A.B., and Bauer, S. (2016). Rapid Determination of Cellulose, Neutral Sugars, and Uronic Acids from Plant Cell Walls by One-step Two-step Hydrolysis and HPAEC-PAD. Bio-protocol 6, e1978. 10.21769/BioProtoc.1978.

76. Foster, C.E., Martin, T.M., and Pauly, M. (2010). Comprehensive compositional analysis of plant cell walls (Lignocellulosic biomass) part I: lignin. J Vis Exp. 10.3791/1745.

77. Patro, R., Duggal, G., Love, M.I., Irizarry, R.A., and Kingsford, C. (2017). Salmon provides fast and bias-aware quantification of transcript expression. Nat Methods 14, 417–419. 10.1038/nmeth.4197.

78. Soneson, C., Love, M.I., and Robinson, M.D. (2015). Differential analyses for RNA-seq: transcript-level estimates improve gene-level inferences. F1000Res 4, 1521. 10.12688/f1000research.7563.2.

79. Robinson, M.D., McCarthy, D.J., and Smyth, G.K. (2010). edgeR: a Bioconductor package for differential expression analysis of digital gene expression data. Bioinformatics 26, 139–140. 10.1093/bioinformatics/btp616.

80. Chen, Y., Lun, A.T., and Smyth, G.K. (2016). From reads to genes to pathways: differential expression analysis of RNA-Seq experiments using Rsubread and the edgeR quasi-likelihood pipeline. F1000Res 5, 1438. 10.12688/f1000research.8987.2.

81. Martin, M. (2011). Cutadapt removes adapter sequences from high-throughput sequencing reads. 2011 *17*, 3. 10.14806/ej.17.1.200.

82. Gan, X., Hay, A., Kwantes, M., Haberer, G., Hallab, A., Ioio, R.D., Hofhuis, H., Pieper, B., Cartolano, M., Neumann, U., et al. (2016). The *Cardamine hirsuta* genome offers insight into the evolution of morphological diversity. Nat Plants 2, 16167. 10.1038/nplants.2016.167.

83. Kim, D., Paggi, J.M., Park, C., Bennett, C., and Salzberg, S.L. (2019). Graph-based genome alignment and genotyping with HISAT2 and HISAT-genotype. Nat Biotechnol 37, 907–915. 10.1038/s41587-019-0201-4.

84. Garcia-Alcalde, F., Okonechnikov, K., Carbonell, J., Cruz, L.M., Gotz, S., Tarazona, S., Dopazo, J., Meyer, T.F., and Conesa, A. (2012). Qualimap: evaluating next-generation sequencing alignment data. Bioinformatics 28, 2678–2679. 10.1093/bioinformatics/bts503.

85. Anders, S., Pyl, P.T., and Huber, W. (2015). HTSeq--a Python framework to work with high-throughput sequencing data. Bioinformatics 31, 166–169. 10.1093/bioinformatics/btu638.

86. Danecek, P., Bonfield, J.K., Liddle, J., Marshall, J., Ohan, V., Pollard, M.O., Whitwham, A., Keane, T., McCarthy, S.A., Davies, R.M., and Li, H. (2021). Twelve years of SAMtools and BCFtools. Gigascience 10. 10.1093/gigascience/giab008.

87. Love, M.I., Huber, W., and Anders, S. (2014). Moderated estimation of fold change and dispersion for RNA-seq data with DESeq2. Genome Biol 15, 550. 10.1186/s13059-014-0550-8.

88. Wu, T., Hu, E., Xu, S., Chen, M., Guo, P., Dai, Z., Feng, T., Zhou, L., Tang, W., Zhan, L., et al. (2021). clusterProfiler 4.0: A universal enrichment tool for interpreting omics data. Innovation (Camb) 2, 100141. 10.1016/j.xinn.2021.100141.

89. Wickham, H. (2016). ggplot2: Elegant Graphics for Data Analysis. Springer-Verlag.

90. Kolde, R. (2019). Pheatmap: pretty heatmaps. R package version 1, 726.

91. Chen, H., and Boutros, P.C. (2011). VennDiagram: a package for the generation of highly-customizable Venn and Euler diagrams in R. BMC Bioinformatics 12, 35. 10.1186/1471-2105-12-35.

92. Aparicio-Puerta, E., Gomez-Martin, C., Giannoukakos, S., Medina, J.M., Scheepbouwer, C., Garcia-Moreno, A., Carmona-Saez, P., Fromm, B., Pegtel, M., Keller, A., et al. (2022). sRNAbench and sRNAtoolbox 2022 update: accurate miRNA and sncRNA profiling for model and non-model organisms. Nucleic Acids Res 50, W710–W717. 10.1093/nar/gkac363.

93. Kumar, S., Stecher, G., Suleski, M., Sanderford, M., Sharma, S., and Tamura, K. (2024). MEGA12: Molecular Evolutionary Genetic Analysis Version 12 for Adaptive and Green Computing. Mol Biol Evol 41. 10.1093/molbev/msae263.

94. Lovell, J.T., Sreedasyam, A., Schranz, M.E., Wilson, M., Carlson, J.W., Harkess, A., Emms, D., Goodstein, D.M., and Schmutz, J. (2022). GENESPACE tracks regions of interest and gene copy number variation across multiple genomes. Elife 11. 10.7554/eLife.78526.

95. McLeay, R.C., and Bailey, T.L. (2010). Motif Enrichment Analysis: a unified framework and an evaluation on ChIP data. BMC Bioinformatics 11, 165. 10.1186/1471-2105-11-165.

96. Brudno, M., Poliakov, A., Minovitsky, S., Ratnere, I., and Dubchak, I. (2007). Multiple whole genome alignments and novel biomedical applications at the VISTA portal. Nucleic Acids Res 35, W669–674. 10.1093/nar/gkm279.

97. R Core Team (2025). R: A Language and Environment for Statistical Computing (R Foundation for Statistical Computing).

98. Posit team (2025). RStudio: Integrated Development Environment for R (Posit Software, PBC).

99. Kassambara, A. (2023). rstatix: Pipe-Friendly Framework for Basic Statistical Tests.

100. Wickham H, F.R., Henry L, Müller K, Vaughan D (2025). dplyr: A Grammar of Data Manipulation.

101. Wickham H, V.D., Girlich M (2025). tidyr: Tidy Messy Data.

102. Rauluseviciute, I., Riudavets-Puig, R., Blanc-Mathieu, R., Castro-Mondragon, J.A., Ferenc, K., Kumar, V., Lemma, R.B., Lucas, J., Cheneby, J., Baranasic, D., et al. (2024). JASPAR 2024: 20th anniversary of the open-access database of transcription factor binding profiles. Nucleic Acids Res 52, D174–D182. 10.1093/nar/gkad1059.

103. Camacho, C., Coulouris, G., Avagyan, V., Ma, N., Papadopoulos, J., Bealer, K., and Madden, T.L. (2009). BLAST+: architecture and applications. BMC Bioinformatics 10, 421. 10.1186/1471-2105-10-421.

104. Edgar, R.C. (2004). MUSCLE: multiple sequence alignment with high accuracy and high throughput. Nucleic Acids Res 32, 1792–1797. 10.1093/nar/gkh340.

105. Waterhouse, A.M., Procter, J.B., Martin, D.M., Clamp, M., and Barton, G.J. (2009). Jalview Version 2--a multiple sequence alignment editor and analysis workbench. Bioinformatics 25, 1189–1191. 10.1093/bioinformatics/btp033.

