## Supplemental Figures S1-S7 for "REVOLUTA regulates cell fate and wall patterning in the fruit endocarp"

### Supplementary figures

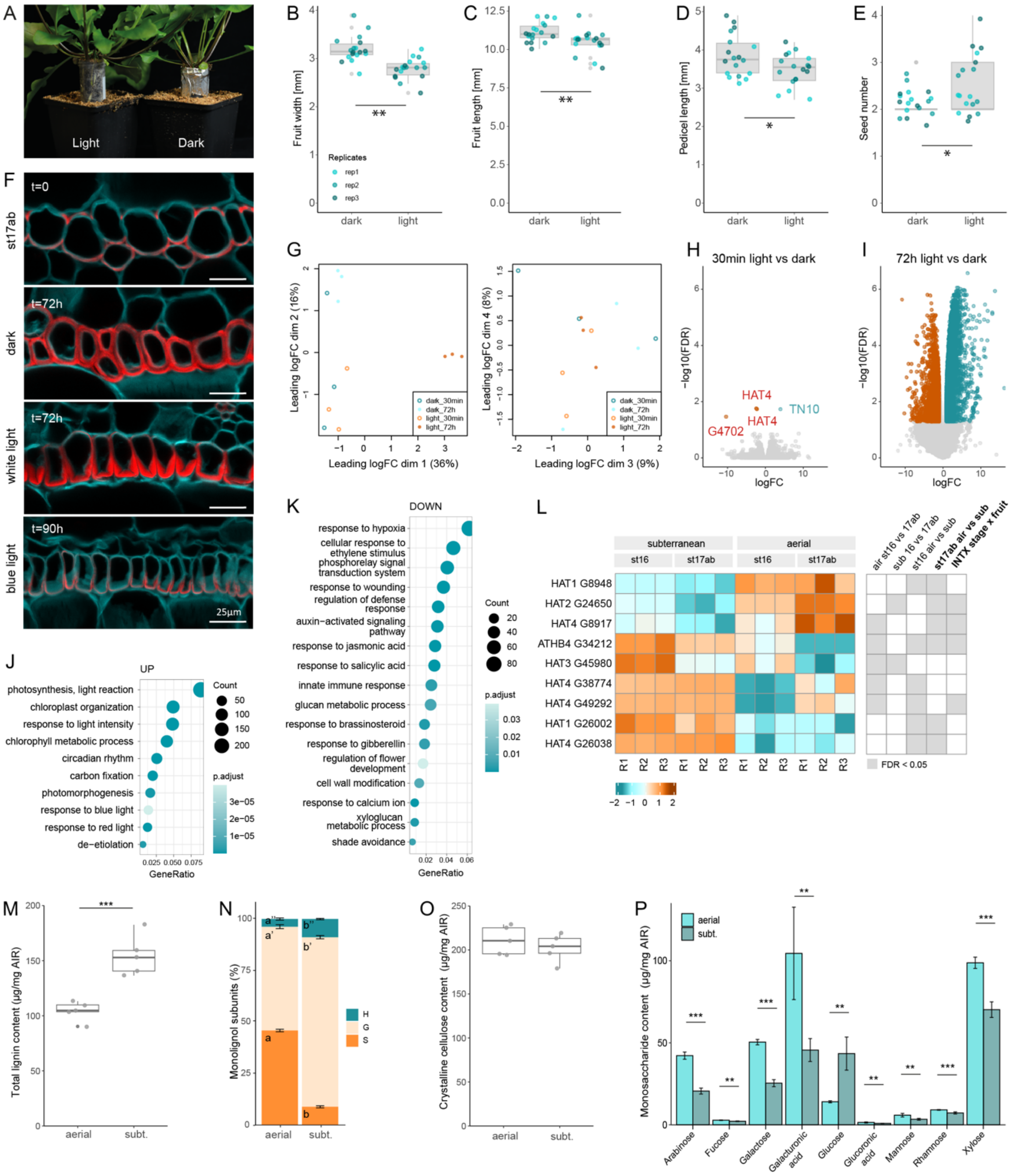


**Figure S1: Light-responsive switch to produce explosive seed pods in *C. chenopodiifolia*, related to Fig. 1. (A)** Experimental setup for light and dark treatments of *C. chenopodiifolia* subterranean fruit. Plant roots and subterranean fruits grew through a soil column within a Falcon tube prior to entering the soil of the black pot. Falcon tubes were fitted with light-proof covers; for light treatment (Light), the cover was removed to expose fruits growing along the transparent tube walls to light; for dark treatment (Dark), the cover remained in place. **(B-E)** *C. chenopodiifolia* subterranean fruit width (B), length (C), pedicel length (D) and seed number (E) after dark or light treatment. Experiments were done in triplicate (n=18). Asterisks denote statistically significant differences at $p < 0.05$ (*) or $p < 0.01$

(**) using Student's t-test or Wilcoxon Mann-Whitney test when the data did not follow linear model assumptions; p = 1.54e-05 (B), 5.76e-03 (C), 3.07e-02 (D), 2.70e-02 (E). **(F)** End*b* of subterranean *C. chenopodiifolia* fruit at stage 17ab (t = 0) and subsequently exposed to dark, white light for 72 h (t = 72 h, ~90 µmol $m^{-2}$ $s^{-1}$) or monochromatic blue light for 90h (t = 90h, ~10 µmol $m^{-2}$ $s^{-1}$). Sections were stained for cellulose (calcofluor white, cyan) and lignin (basic fuchsin, red) (n=6 for t=0h and dark 72h, n=9 for light 72h, n=10 for blue light 90h). **(G)** Principal component analysis of RNA-seq data from *C. chenopodiifolia* subterranean fruit valve samples exposed to light or dark. **(H-I)** Volcano plots of DEGs (logFC ≥|1|, FDR < 0.05) for light vs dark comparisons at 30 min (H) and 72 h (I). All four DEGs are annotated for the 30-min time point. **(J-K)** Selected GO terms enriched in the 3232 up-regulated (J) and 1846 down-regulated (K) DEGs in the comparison of light and dark treatment at 72-h time point. **(L)** HD-ZIPII genes shown in main Fig. 1I in aerial vs subterranean fruit valves at stages 16 and 17ab and the interaction of fruit type and fruit stage (INTX stag x fruit). Grey boxes indicate statistically significant differences (logFC ≥ |1|, FDR < 0.05) and heat map shows differential gene expression as logFC. **(M-P)** Boxplots and barplots of lignin, monolignol subunits, crystalline cellulose, and matrix monosaccharides in mature aerial vs subterranean (subt.) fruit valves, shown as µg per mg of alcohol-insoluble residue (AIR) for acetylbromide-soluble lignin (M), glucose (O) and matrix polymer sugars (P) or percentage of monolignol subunit (H-, G- or S-type) (N). *** denotes statistical significance at p < 0.001, ** at p < 0.01 using Student's *t*-test or Wilcoxon's test when normality conditions are not fulfilled. Different letters indicate significant differences between fruit type (p < 0.05). Scale bar: 25 µm (F).

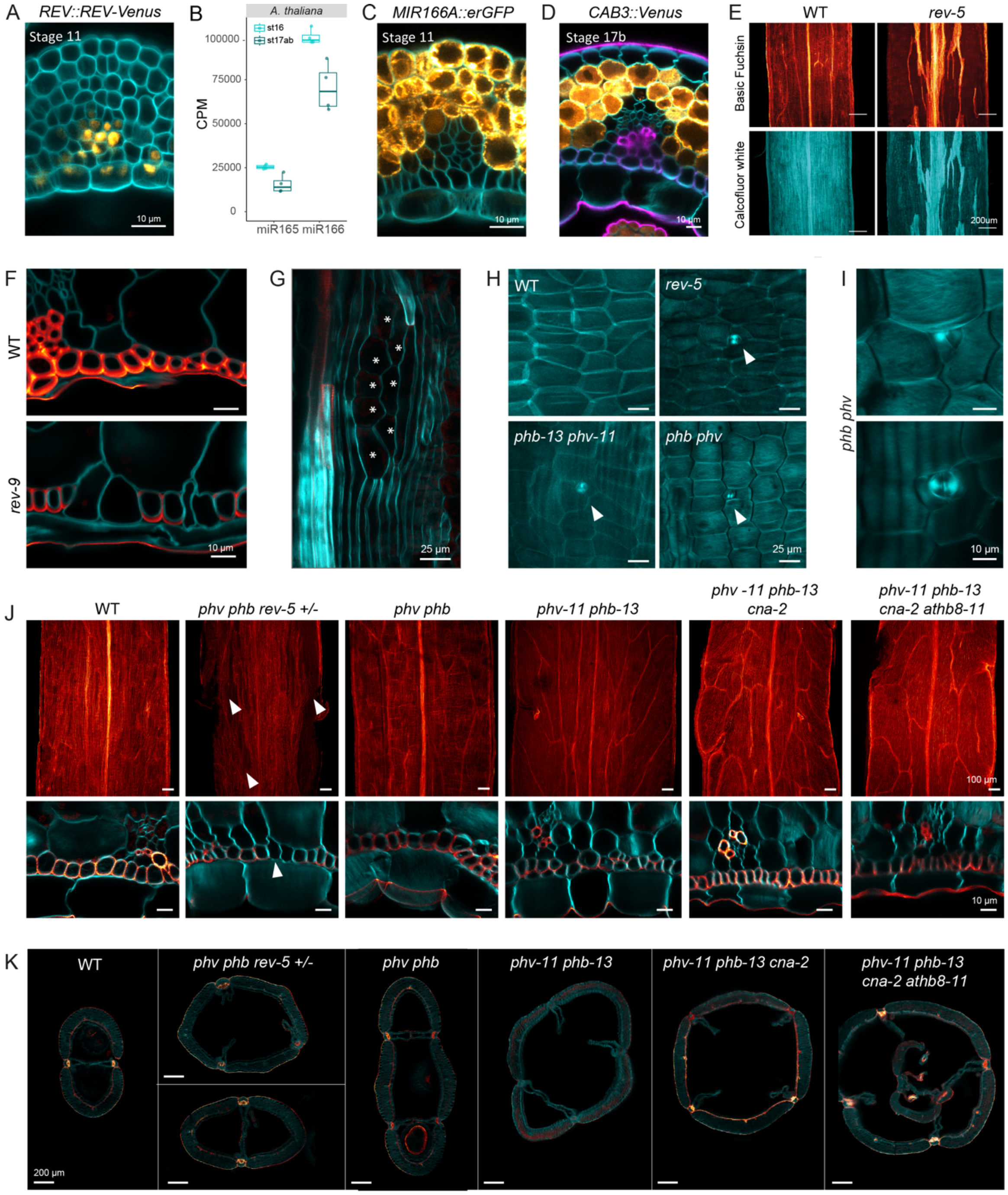


**Figure S2: *REVOLUTA* regulates endocarp *b* identity in Arabidopsis, related to Fig. 2. (A)** *REV::REV-mVenus* signal (Orange Hot LUT) and cellulose stained with calcofluor white (cyan) in transverse section of an Arabidopsis stage 11 fruit valve (n=8). **(B)** Mature miR165 and miR166 abundance in counts per million (CPM) obtained from miRNA sequencing of Arabidopsis fruit valves at stage 16 and 17ab. **(C)** *MIR166A::erGFP* expression (Orange Hot LUT) and cellulose stained with calcofluor white (cyan) in fruit transverse sections of Arabidopsis stage 11 fruit valve (n=3). **(D)** *CAB3::3xVenus-RCl2A* expression (Orange Hot LUT) in transverse section of an Arabidopsis stage 17b fruit valve stained for cellulose (calcofluor white, cyan) and lignin (basic fuchsin, magenta) (n=4). **(E)** Arabidopsis wild-type (WT) and *rev-5* mutant fruit valves at stage 17b stained for cellulose (cyan) and lignin (Red Hot LUT) (n = 21 WT, 22 *rev-5*). Maximum projection of adaxial surface of end*b* cell layer. **(F-K)** Arabidopsis fruit valves stained for cellulose (calcofluor white, cyan) and lignin (basic fuchsin, Red Hot LUT). (F) Transverse sections through end*b* cells of wild-type (WT) and *rev-9* (total n=6). (G) Maximum projection of adaxial end*b* surface in *rev-5*; asterisks indicate mesocarp-like cells (n=22). (H) Maximum projection of adaxial endocarp *a* surface of WT (n=18), *rev-5* (n=22), *phv-11 phb-13* (n=3) and *phb phv* (n=4) mutants. Arrowheads indicate ectopic stomata. (I) Close up view of ectopic stomata in *phb phv* (n=4); images show two different planes through the same z-stack. (J-K) Maximum projection of adaxial end*b* surface (upper panels, J) and transverse fruit sections (lower panels, J, K) of Arabidopsis WT (n=18) and *HD-ZIPIII* multiple mutants: segregating *phv phb rev-5 +/-* (n=9), *phv phb* (n=4), *phv-11 phb-13* (n=3), *phv-11 phb-13 cna-2* (n=8) and *phv-11 phb-13 cna-2 athb8-1* (n=5). WT panel reused from main Fig. 2D Scale bars: 10 µm (A, C, D, F, I, J), 25 µm (G, H), 100 µm (upper panels, J), 200 µm (E, K).

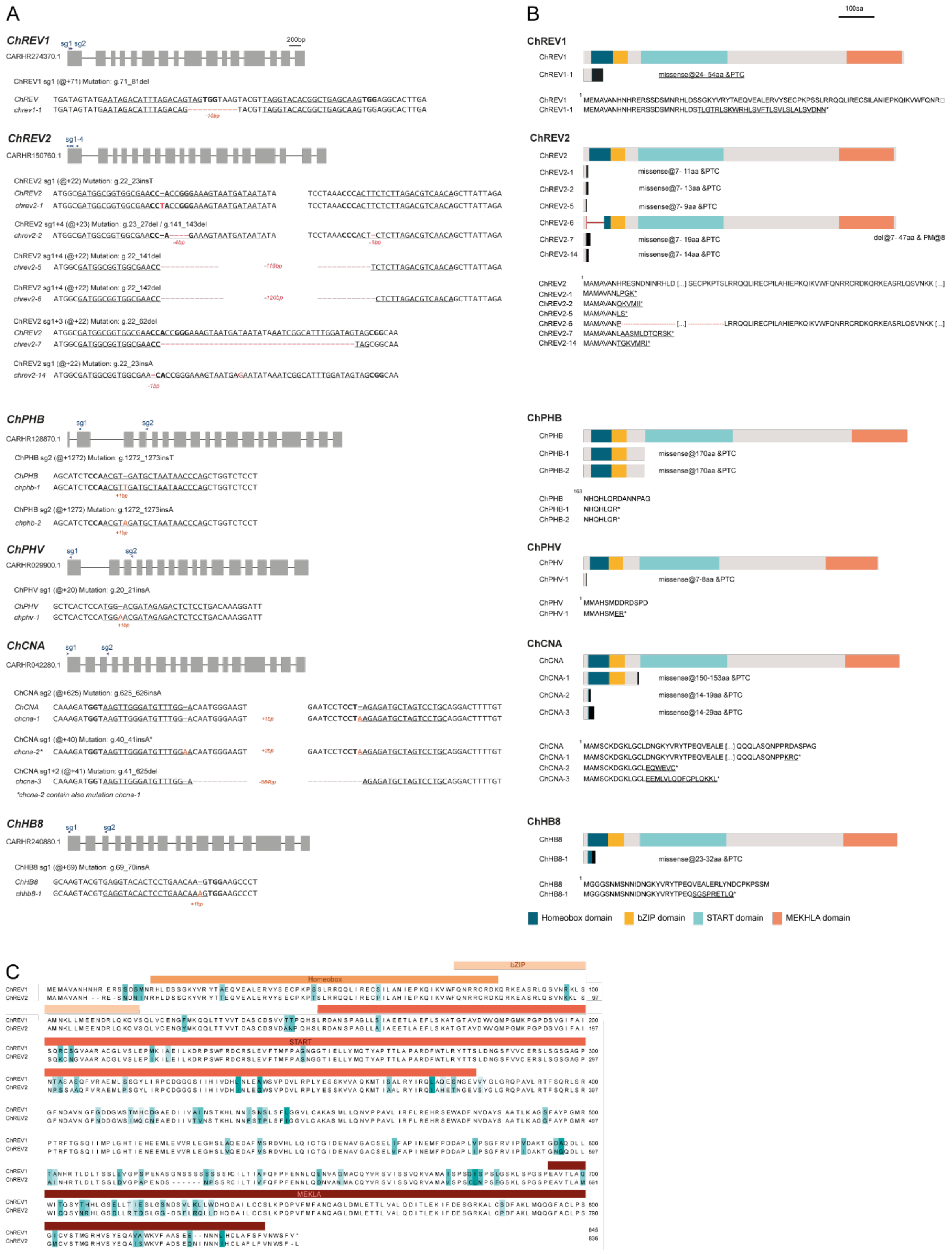

**Figure S3: Characterization of *C. hirsuta* CRISPR/Cas9 *HD-ZIPIII* alleles, related to Fig. 3. (A)** Schematic of *C. hirsuta ChREV1, ChREV2, ChPHB, ChPHV, ChCNA* and *ChHB8* genes, CRISPR sgRNAs and alleles generated in this study. Gray boxes: coding DNA sequences, black lines: introns, blue arrows: sequences targeted by CRISPR sgRNAs. @+ symbol on WT sequence indicates position of mutation counting from the start codon. For example, g.71_81del indicates a deletion of ten nucleotides. Mutations are indicated in orange. Underlined sequence: CRISPR sgRNA, bold text: PAM sequence. **(B)** Putative translational products of CRISPR/Cas9 HD-ZIPIII alleles. (*Top*) Schematic of the proteins encoded by WT and mutant alleles. Dark blue box: homeobox domain, orange box: bZIP domain, light blue box: START domain, yellow box: MEKHLA domain. Black rectangles indicate putative peptide from missense translation in mutant alleles. Mutant alleles are represented up to the premature termination codon (PTC). Description of each mutant protein, indicating the number of amino acid (aa) residues resulting from missense translation and the PTC. For example, "missense@24-54aa & PTC" indicates that aa residues 24 to 54 result from missense translation, followed by a PTC. (*Bottom*) Predicted peptide sequences encoded by WT and mutant alleles. Underlined characters indicate aa residues that result from missense translation. **(C)** Alignment of *C. hirsuta* ChREV1 and ChREV2 protein sequences using MUSCLE. Scale bars: 200 bp (A), 100 aa (B).

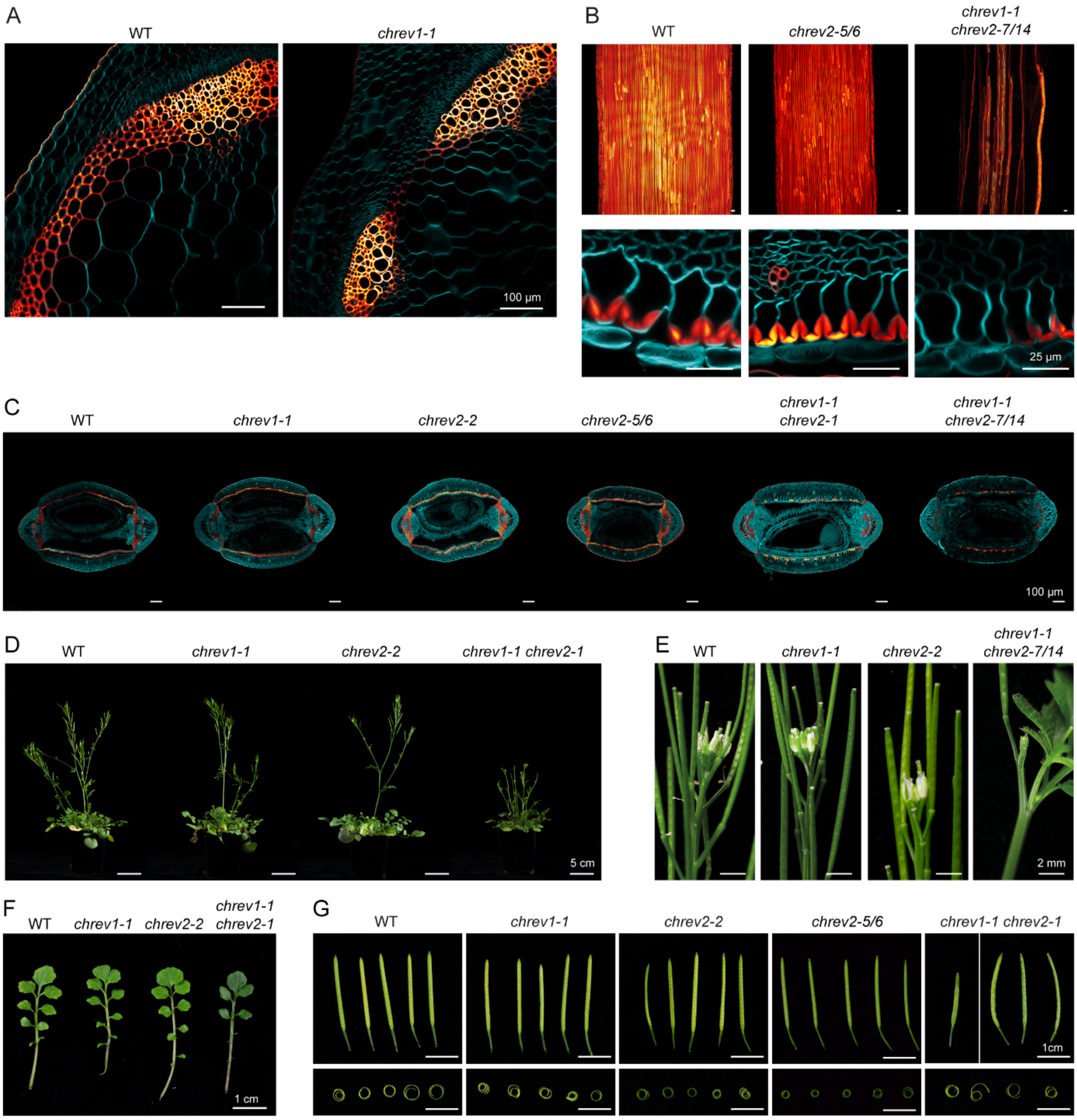


**Figure S4: Phenotypic analysis of *C. hirsuta rev1, rev2* single and double mutants, related to Fig.3.** *C. hirsuta* genotypes: WT (n=23), *chrev1-1* (n=18), *chrev2-2* (n=4), *chrev2-5/6* (T1, n=1), *chrev1-1 chrev2-1* (n=8), *chrev1-1 chrev2-7/14* (T1, n=1). **(A-C)** Stem transverse sections (A) and stage 17b fruit (B-C) stained for cellulose (calcofluor white, cyan) and lignin (basic fuchsin, Red Hot LUT) shown as maximum projections of the adaxial end*b* surface (upper panels, B), end*b* cells in transverse section (lower panels, B) and fruit transverse sections (C). **(D)** Plants at 7.5 weeks after germination. **(E)** Main inflorescence at 6.5 weeks. **(F)** Last rosette leaf. **(G)** Whole fruit and removed valves (n=4-5). Scale bars: 100 µm (A, C), 25 µm (B), 5 cm (D), 2 mm (E), 1 cm (F-G).

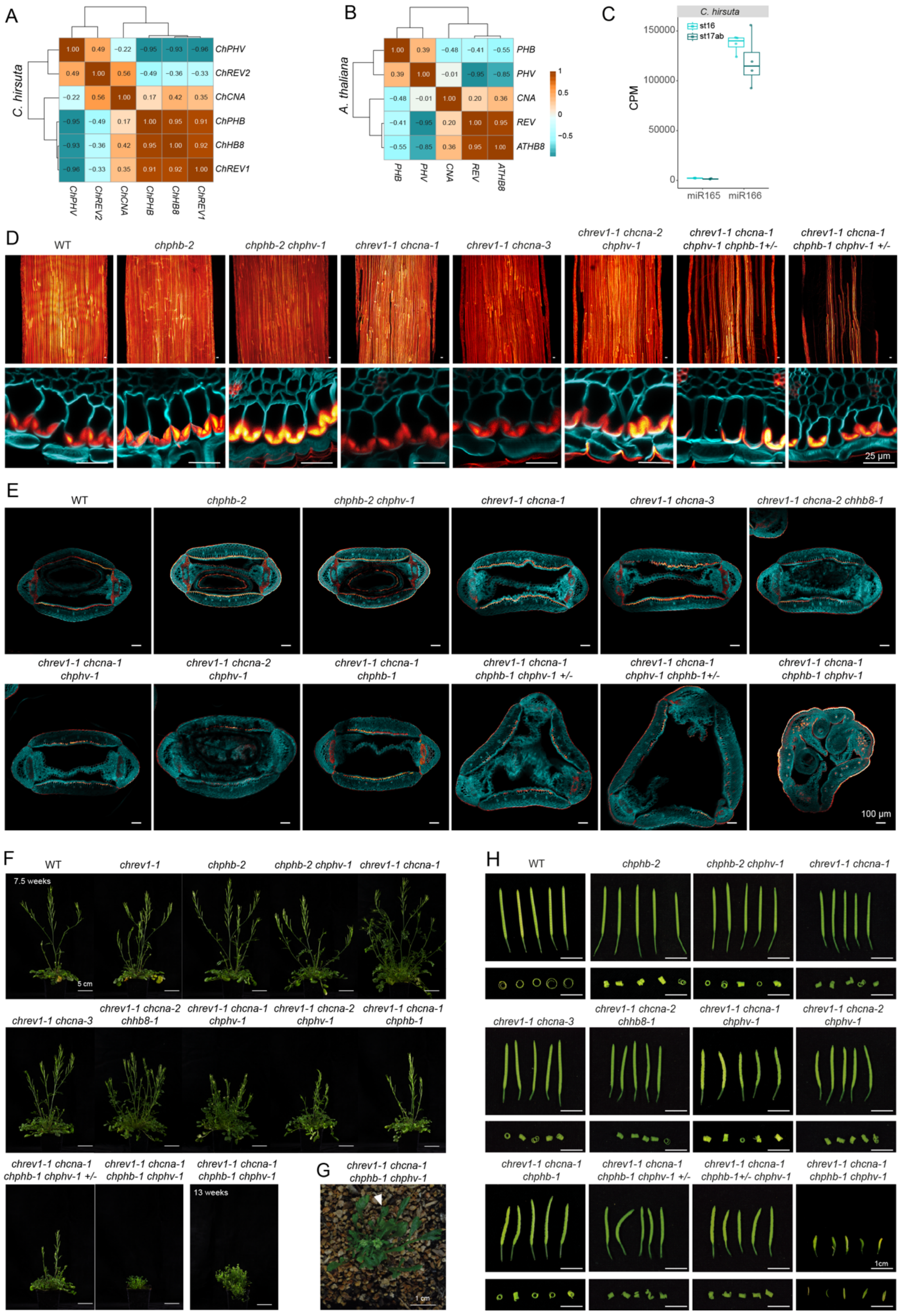


**Figure S5: Phenotypic analysis of multiple *HD-ZIPIII* mutants in *C. hirsuta*, related to Fig. 3. (A-B)** Pearson correlation coefficient matrix of *HD-ZIPIII* gene expression from RNA-seq of *C. hirsuta* (A) and Arabidopsis (B) fruit valves at stage 16 and 17ab. **(C)** Mature miR165 and miR166 abundance in counts per million (CPM) obtained from miRNA sequencing of *C. hirsuta* fruit valves at stage 16 and 17ab. **(D-H)** *C. hirsuta* WT (n=23: D-F,H) and the following *HD-ZIPIII* mutants: *chrev1-1* (n=18: F), *chphb-2* (n=8: D-F, H)*, chphb-2 chphv-1* (n=10: D-F, H)*, chrev1-1 chcna-1* (n=11: D-F, H)*, chrev1-1 chcna-3* (n=14: D-F, H)*, chrev1-1 chcna-2 chhb8-1* (n=10: D-F, H)*, chrev1-1 chcna-1 chphv-1* (n=10: E-F,H)*, chrev1-1 chcna-2 chphv-1* (n=10: D-F, H)*, chrev1-1 chcna-1 chphb-1* (n=8: E-F,H)*,* segregating *chrev1-1 chcna-1 chphv-1 chphb-1 +/-* (n=4: E-F,H), *chrev1-1 chcna-1 chphb-1 chphv-1 +/-* (n=14: D-F, H) and *chrev1-1 chcna-1 chphb-1 chphv-1* (n=6: E-F,H). Stage 17b fruit stained for cellulose (calcofluor white, cyan) and lignin (basic fuchsin, Red Hot LUT). Maximum projection of adaxial end*b* surface (upper panels, D), end*b* cells in transverse section (lower panels, D) and fruit transverse sections (E). Plants at 7.5 or 13 weeks after germination (F), *chrev1-1 chcan-1 chphb-1 chhv-1* quadruple mutant plant at 6 weeks after germination; arrow indicates abaxialised trumpet-shaped leaf (G). Whole fruit and removed valves (H, n =4-5). Scale bars: 25 µm (D), 100 µm (E), 5 cm (F), 1 cm (G-H). WT images reused from Fig. S4B-D, G.

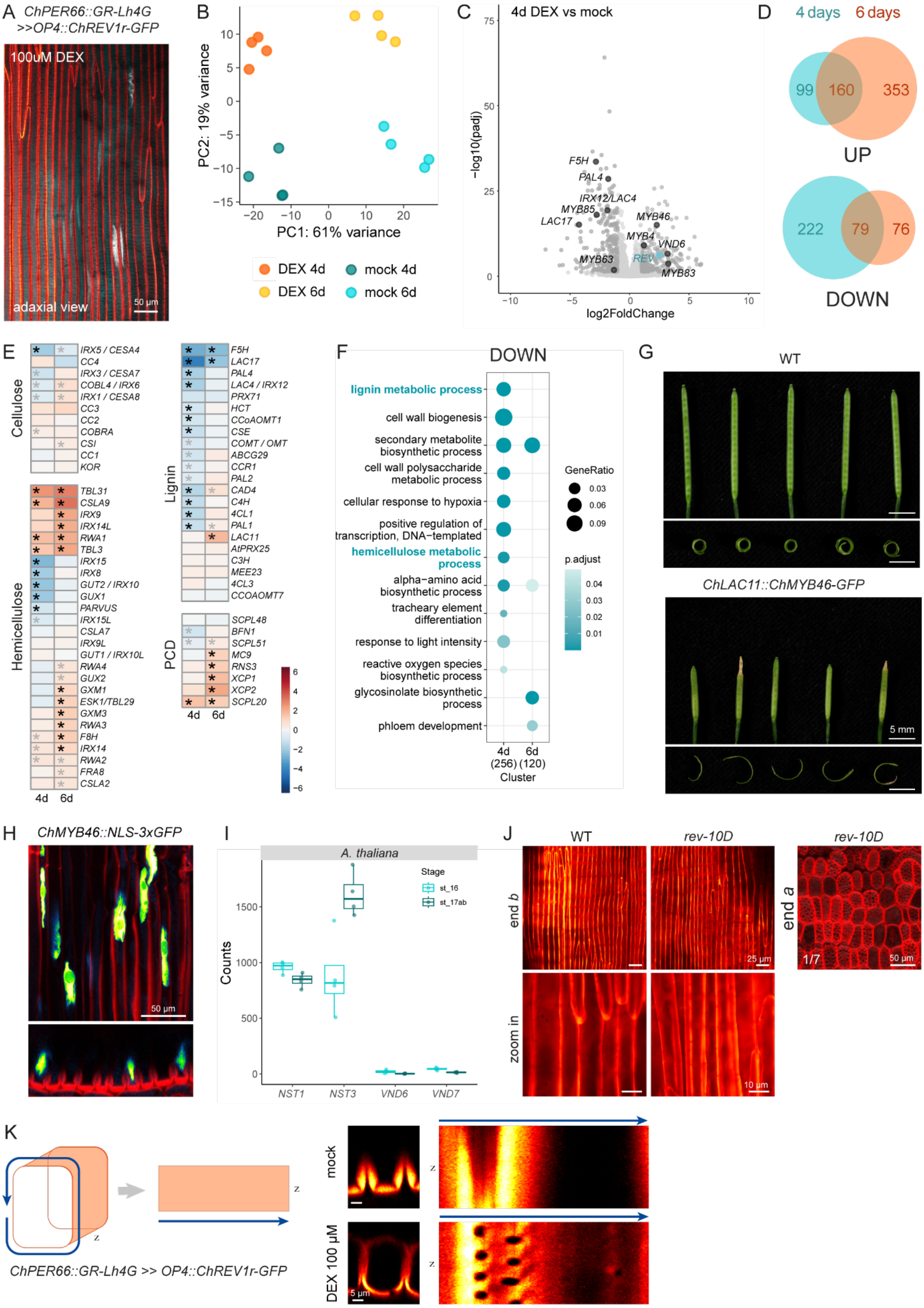


**Figure S6: *REVOLUTA* is sufficient to switch endocarp *b* SCW pattern in *C. hirsuta*, related to Fig. 4.**

**(A)** Maximum projection of adaxial end*b* surface of *ChPER66::GR-Lh4G>>pOP4::ChREV1r-GFP* (Grey

LUT) treated with 100 µM DEX showing ChREV1r-GFP signal in end*b* nuclei. Lignin stained with basic fuchsin (red) (n=9). **(B)** Principal component analysis of RNA-seq data from *C. hirsuta ChPER66::GR-Lh4G>>pOP4::ChREV1r-GFP* fruit valves treated with mock or 100 µM DEX. **(C-D)** Volcano plot (C) and Venn diagram (D) of up- and down-regulated differentially expressed genes (DEGs), (logFC ≥ |1|, p-adj < 0.05) between mock and DEX-treated *C. hirsuta ChPER66::GR-Lh4G>>pOP4::ChREV1r-GFP* fruit valves after 4-day (C) or 4- and 6-day (D) treatments. **(E)** Heat map of log fold-change (logFC) in the DEX vs mock comparison at 4 days and 6 days for genes involved in cellulose, hemicellulose, lignin biosynthesis and programmed cell death (PCD). Black stars indicate significant values with logFC ≥ |1|, p-adj < 0.05, grey star: logFC < |1|, p-adj < 0.05. **(F)** Selected GO terms enriched in the 256 and 120 down-regulated DEGs for the DEX vs mock comparison at 4 days and 6 days. **(G)** Whole fruit and removed valves of WT and *ChLAC11::ChMYB46-GFP* (n=12). **(H)** Maximum projection (upper panel) and optical section (lower panel) of *ChMYB46::NLS-3xGFP* signal (Green Blue Fire LUT) in end*b* cells (n=3). **(I)** Relative counts for Arabidopsis *NST1/3* and *VND6/7* genes from RNA-seq of fruit valves at stage 16 and 17ab. **(J)** Maximum projection of adaxial end*b* surface of Arabidopsis wild type and dominant *rev-10D* mutant stained for lignin (basic fuchsin, Red Hot LUT). One out of seven samples showed ectopic lignification of the endocarp *a* layer in a pitted pattern (n=7). **(K)** Optical section (left) obtained from z-stack of end*b* cells stained for lignin (basic fuchsin, Red Hot LUT) in *ChPER66::GR-Lh4G> pOP4::ChREV1r-GFP* fruit treated for 7d with mock or 100 µM DEX. Kymographs (right) obtained by measuring basic fuchsin signal along a line following the cell outline, as depicted in the small cartoon above (n=3). Scale bar: 50 µm (A, H), 5 mm (G), 25 µm (J, upper panels), 10 µm (J, lower panels), 50 µm (J, right panel), 5 µm (K).

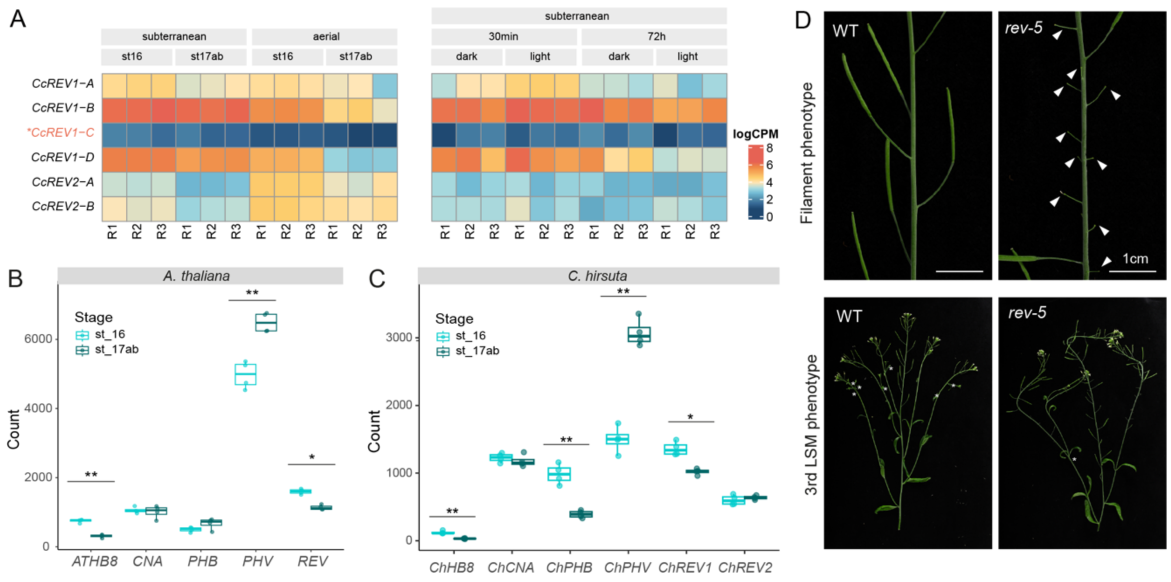


**Figure S7: *REVOLUTA* gene duplication and divergence in *C. chenopodiifolia,* related to Fig. 5. (A)** Heat map of log Count Per Million (logCPM) values for six *C. chenopodiifolia REV* genes (*CcREV1-A/B/C/D* and *CcREV2-A/B*) in comparisons between subterranean and aerial fruit valves at stages 16 and 17ab *(left panel)* and subterranean fruit valves exposed to dark vs light for 30 min and 72 h *(right panel).* * indicates that *CcREV1-C* is a likely pseudogene. Values are not normalized in order to visualize low counts for *CcREV1-C.* **(B-C)** Relative counts for *HD-ZIPIII* genes obtained from RNA-seq of Arabidopsis (C) and *C. hirsuta* (D) fruit valves at stages 16 and 17ab. Stars indicate significant differences (**logFC ≥ |1|, adjusted p-value (padj) < 0.05; *logFC < |1|, padj < 0.05). **(D)** Arabidopsis *rev-5* phenotypes: defects in flower and axillary meristem formation. Filamentous structures and carpel-less flowers form in *rev-5* (arrows), compared to normal fruit in WT (*upper panels).* Tertiary lateral shoot meristems (3$^{rd}$ LSM) formed in the axils of cauline leaves in WT secondary branches (arrows) are reduced in *rev-5* (*lower panels)* (n=26). Scale bar: 1 cm (D).
